# A biomimetic five-module chimeric antigen receptor (^5M^CAR) designed to target and eliminate antigen-specific T cells

**DOI:** 10.1101/2020.01.24.916932

**Authors:** Shio Kobayashi, Martin A. Thelin, Heather L. Parrish, Neha R. Deshpande, Mark S. Lee, Alborz Karimzadeh, Monika A. Niewczas, Thomas Serwold, Michael S. Kuhns

## Abstract

T cells express clonotypic T cell receptors (TCRs) that recognize peptide antigens in the context of class I or II MHC molecules (pMHCI/II). These receptor modules associate with three signaling modules (CD3γε, δε, and ζζ), and work in concert with a coreceptor module (either CD8 or CD4), to drive T cell activation in response to pMHCI/II. Here we describe a first generation biomimetic 5-module chimeric antigen receptor (^5M^CAR). We show that: (i) chimeric receptor modules built with the ectodomains of pMHCII assemble with CD3 signaling modules into complexes that redirect cytotoxic T lymphocyte (CTL) specificity and function in response to the clonotypic TCRs of pMHCII-specific CD4^+^ T cells; and, (ii) surrogate coreceptor modules enhance the function of these complexes. Furthermore, we demonstrate that adoptively transferred ^5M^CAR-CTLs can mitigate type I diabetes by targeting autoimmune CD4^+^ T cells in NOD mice. This work provides a framework for the construction of biomimetic ^5M^CARs that can be used as tools to study the impact of particular antigen-specific T cells in immune responses, and may hold potential for ameliorating diseases mediated by pathogenic T cells.

## Main

T cells scan major histocompatibility complex (MHC) molecules on the surfaces of cells for the presence of peptide antigens (pMHC) derived from microbes, vaccines, or tumor cells with their clonotypic T cell receptors (TCRs). If the dwell time of the TCR on the pMHC is of sufficient duration, a T cell will become activated and differentiate to helper (Th), cytotoxic (CTL), regulatory (Treg), or memory (Tm) T cells that are essential for long-lived immunity^1,2^. CD4^+^ Th subsets provide help for effective CTL, B cell, and innate immune cell function upon immunologic challenge, Tregs are crucial for peripheral tolerance to self and commensal antigens, Tm allow for rapid antigen-specific responses, and CTLs can eliminate infected or cancerous cells^3–6^. However, the activity of each T cell subset can be counter-productive if conditions are such that they result in the induction of allergies, asthma, autoimmunity, transplant rejection, or, in the case of Tregs, the protection of tumors. Considerable effort has thus been focused on developing strategies to: determine how T cells of a particular pMHC-specificity impact an immune response; enhance T cell responses to fight infections or tumors; or, mitigate T cell-mediated pathologies.

Chimeric antigen receptors (CARs) have gained attention as a technology that can redirect T cell specificity and function for novel purposes (**Fig. 1a**). The archetypal CAR design consists of a single-chain module (referred to here as ^1M^CARs) wherein ligand specificity (e.g. tumor antigens) is usually conferred via an antibody-derived Fv, while intracellular signaling is directed through a tandem array of known signaling motifs^7,8^. Work to improve ^1M^CAR efficacy has resulted in the development and testing of numerous variations on the initial design, including fragmented domains that form ^1M^CARs upon final assembly^8,9^. Yet, despite these efforts, ^1M^CARs require ~100-1000 fold more antigen to direct CTL responses than their natural counterpart^9–11^. Because the basic template of these *de novo* receptors was established before we better appreciated how natural receptors trigger, ^1M^CARs may lack the means to efficiently relay ligand-specific information across the cell membrane; thus, there may be practical limits to what can be achieved with variants of the archetypal ^1M^CAR design^8,9,12^.

**Fig. 1:**
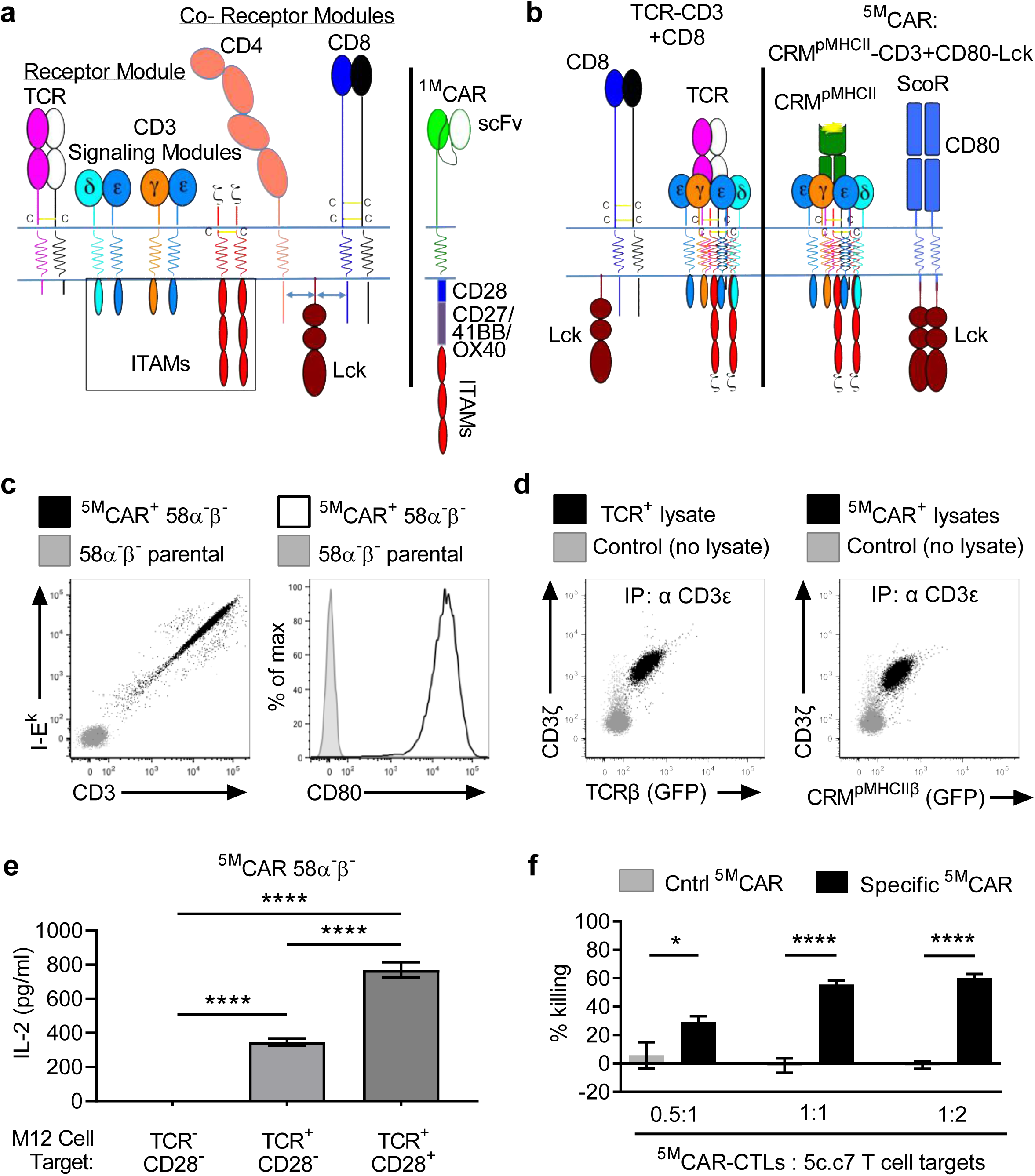
Structure, assembly, and function of biomimetic ^5M^CARs. **a**, The 5 modules that drive pMHC-specific T cell activation (TCR, CD3*γε*, CD3*δε*, CD3*ζζ*, and CD4/CD8) are illustrated in comparison with a 3^rd^ generation single-module CAR (^1M^CAR). **b**, Illustration of the TCR-CD3 complex and CD8 in comparison with the CRM^pMHCII^-CD3 complex and CD80-Lck ScoR of ^5M^CARs. **c**, Flow cytometry plots showing I-E^k^, CD3, and CD80 expression on parental 58α^−^β^−^ cells and MCC:I-E^k+ 5M^CAR-58α^−^β^−^ cells. **d**, FFLISA of TCR-CD3 and CRM^pMHCII^-CD3 complexes. Anti-CD3ε beads incubated without lysate (grey), or with lysates from TCR-CD3^+^ 58α^−^β^−^ cells or MCC:I-E^k+ 5M^CAR-58α^−^β^−^ cells (black), were analyzed by flow cytometry for TCRβ or CRM^pMHCII^^β^ (GFP) and CD3*ζ* association. **e**, IL-2 production by MCC:I-E^k+ 5M^CAR-58α^−^β^−^ cells after 16hr coculture with parental M12 B cells (TCR^−^, CD28^−^), 2B4 TCR^+^ M12 cells, or 2B4 TCR^+^ CD28^+^ M12 cells was quantified by ELISA (****p < 0.0001, one-way ANOVA with Tukey’s posttest). **f**, ^5M^CAR-CTL killing of CD4^+^ 5c.c7 TCR Tg T cell targets. Percent killing of targets co-cultured with control (Hb:I-E^k^) or specific (MCC:I-E^k^) ^5M^CAR-CTLs was measured by flow cytometry and is presented relative to number of targets cultured in the absence of ^5M^CAR-CTLs (*p < 0.05, ***p < 0.001, ****p < 0.0001 by unpaired, two-tailed *t*-test). All data are representative of at least 2 experiments.

An alternative approach is to employ biomimetic engineering to develop CARs that mirror the operating principles of the highly sensitive and specific 5-module receptors that have evolved to drive T cell response to pMHC. In brief, the TCR is the receptor module (module 1). It binds pMHC and relays information to the immunoreceptor tyrosine-based activation motifs (ITAMs) of the three associated signaling modules (CD3γε, δε, and ζζ; modules 2-4)^12,13^. CD4 and CD8 are coreceptors that represent the fifth module on CD4^+^ or CD8^+^ T cells; they bind MHCII or MHCI, respectively, and associate non-covalently with the Src kinase, p56^Lck^ (Lck), that phosphorylates CD3 ITAMs. The coreceptors sequester Lck away from TCR-CD3 complexes until either CD4 or CD8 and the TCR both bind pMHC, at which point Lck is positioned proximal to the CD3 ITAMs to initiate signaling^14–17^. These 5-module pMHC-receptors can signal in response to a single agonist pMHC, direct CTL killing against just three pMHC, and direct distinct T cell responses according to the quantity and quality of the pMHC^1,2,11,18–23^. Given the extraordinary sensitivity, specificity, and cell fate-directing ability of the natural pMHC receptors, engineering biomimetic versions of multi-module CARs could expand and enhance the applications of CAR-T cell therapy.

To this end we engineered a 5-module chimeric antigen receptor (^5M^CAR) system based on the operating principles of the multi-module receptors that have evolved to direct T cell responses to pMHC. Specifically, we designed a Chimeric Receptor Module (CRM) built with the ectodomain of pMHCII (CRM^pMHCII^) to associate with the three CD3 signaling modules. This design maintained the natural receptor:ligand binding kinetics that drive T cell activation, and enabled us to redirect CTL specificity against clonotypic TCRs expressed by pathogenic T cells. We also engineered a surrogate coreceptor (ScoR) composed of CD80 fused to Lck as the fifth module of our ^5M^CAR. We report that ^5M^CAR-T cell hybridomas make IL-2 in a ligand-specific (i.e. TCR-specific) manner, that ^5M^CAR-CTLs can kill CD4^+^ T cells both *in vitro* and *in vivo* in a TCR-specific manner, and that ^5M^CAR-CTLs targeting autoimmune CD4^+^ T cells can prevent disease in non-obese diabetic (NOD) mouse models. Our results demonstrate that biomimetic ^5M^CARs can redirect CTL specificity and function for novel applications. They also provide data suggesting that the current design could be used therapeutically to mitigate diseases caused by pathogenic T cells.

## Results

### Design and *in vitro* characterization of a 1^st^ generation ^5M^CAR

When designing our 1^st^ generation ^5M^CAR, we decided to maintain pMHCII-TCR interactions as the core receptor-ligand recognition event. Doing so allowed us to preserve the key biophysical properties that have evolved to mediate antigen recognition. Using this strategy, we generated a ^5M^CAR system with the potential to redirect CTLs to target and kill CD4^+^ T cells via recognition of their clonotypic TCRs.

First, we engineered a pMHCII-based chimeric receptor module (CRM^pMHCII^, module 1) to assemble with the CD3*γε*, *δε*, and *ζζ* signaling modules (modules 2-4) into a functional complex (**Fig. 1b and sFig. 1**). The basic design for all CRM^pMHCII^ modules used herein involved fusing the MHCIIα and MHCIIβ ectodomains (ECDs) to the connecting peptides (CP), transmembrane domains (TMD), and intracellular domains (ICD) of the TCRα and TCRβ subunits, respectively. The peptide antigen was N-terminally tethered to the MHCIIβ region to ensure expression of a single CRM^pMHCII^ species, while mEGFP was tethered at the C-terminus to aid in detection. We expected the CRM^pMHCII^ to assemble with the CD3*γε*, *δε*, and *ζζ* modules via interactions in the CP and TMD, and provide specificity for cognate TCRs.

We also engineered a surrogate coreceptor (ScoR) composed of the CD80 ECD and TMD fused to Lck (module 5). CD80 naturally interacts with CD28 on naïve T cells as well as CTLA-4 on antigen experienced T cells. The underlying logic for this design is that because CD28, CTLA-4, and the TCR are reported to inhabit similar microdomains early during immunological synapse formation, a CD80-Lck fusion would bind its pairing partners and localize Lck in proximity to the pMHCR-CD3 complex (**Fig. 1b and sFig.1**)^24,25^.

To evaluate the expression and function of our prototype ^5M^CAR components we retrovirally transduced 58α^−^β^−^ cells (a T cell hybridoma line that lacks TCRα and TCRβ^26^) to express both the CRM^pMHCII^ and the CD80-Lck ScoR. The CRM^pMHCII^ was built using the murine MHCII I-E^k^ presenting the model peptide moth cytochrome C (MCC88-103; MCC:I-E^K^). Flow cytometry analysis revealed cell surface expression of MCC:I-E^k^, CD3ε, and CD80 on the transduced but not parental 58α^−^β^−^ cells (**Fig 1c**). The proportional expression of MCC:I-E^k^ and CD3ε suggested that the CRM^pMHCII^ module was assembling with the CD3 modules.

We also performed a flow cytometry-based fluorophore-linked immunosorbant assay (FFLISA) of detergent lysates from TCR^+^ and ^5M^CAR^+^ 58α^−^β^−^ cells to confirm that TCR-CD3 and CRM^pMHCII^-CD3 complexes assemble analogously^14^. Latex beads coated with anti-CD3ε monoclonal antibodies (mAbs) were incubated with lysates and then stained with anti-CD3*ζ* antibodies for analysis by flow cytometry. The TCR-CD3^+^ and CRM^pMHCII^-CD3^+^ samples showed similar levels of GFP and CD3*ζ* signal (**Fig. 1d**), demonstrating that the TCRβ-GFP and MHCIIβ-GFP co-immunoprecipitate (IP) with both CD3ε and CD3*ζ* subunits at similar levels.

To evaluate if our ^5M^CAR^+^ 58α^−^β^−^ cells can respond to cells expressing a specific TCR, we measured IL-2 production after 16 hours of coculture with: parental M12 cells that are TCR and CD28 negative; M12 cells transduced to express TCR-CD3 complexes; or, M12 cells transduced to express both TCR-CD3 complexes and CD28^27^. The 2B4 TCR that binds MCC:I-E^k^ was expressed on the M12 target cells^28^. No IL-2 was produced in response to the parental M12 cells, IL-2 was produced in response to TCR^+^ M12 cells, and ~2x more IL-2 was produced in response to TCR^+^CD28^+^ M12 cells (**Fig. 1e**). Because M12 cells are a B cell lymphoma line and do not make IL-2, the IL-2 measured in this assay was produced by the ^5M^CAR^+^ 58α^−^β^−^ cells. These data establish that our MCC:I-E^k^-based ^5M^CAR system can direct a TCR-specific response.

Next we asked if ^5M^CARs can redirect CTLs to kill CD4^+^ T cells expressing the 5c.c7 TCR, which also recognizes MCC:I-E^k^ as an agonist pMHCII^28^. CD8^+^ T cells from B10.A mice were activated *in vitro* and transduced to express the ^5M^CAR components. ^5M^CAR-CTLs expressing a MCC:I-E^k^-based CRM^pMHCII^ were generated to specifically target 5c.c7 CD4^+^ T cells, while ^5M^CAR-CTLs expressing an Hb:I-E^k^-based CRM^pMHCII^ were generated as negative controls that should not target 5c.c7 CD4^+^ T cells^15,29^. Target CD4^+^ T cells from 5c.c7 TCR transgenic mice were either cultured alone, or with the specific and control ^5M^CAR-CTLs at varying ratios, to assess killing. The number of target CD4^+^ 5c.c7 T cells that remained after 16hrs culture in the absence of ^5M^CAR-CTLs was enumerated by flow cytometry in order to establish a no-killing baseline that was then used to calculate the percent killing by the ^5M^CAR-CTLs. We observed significantly higher killing of target T cells by the specific ^5M^CAR-CTLs when compared with the control ^5M^CAR-CTLs at all effector:target ratios (**Fig. 1f**). No off-target killing was observed in the control group. These data demonstrate that ^5M^CAR-CTLs can kill CD4^+^ T cell targets *in vitro* in a CRM^pMHCII^-dependent manner.

### Targeting pathogenic CD4^+^ T cells with ^5M^CAR-CTLs *in vitro*

Having established the basic functionality of our ^5M^CAR, we asked if we could use it to redirect CTLs to kill CD4^+^ T cells that are reactive to a self-pMHCII. Here we used CD4^+^ T cells expressing the BDC2.5 TCR as targets because: 1) CD4^+^ T cells from BDC2.5 TCR transgenic (Tg) mice recognize a self-peptide antigen derived from pancreatic β-cells presented on the murine MHCII I-A^g7^; and, 2) they can mediate β-cell destruction and type-1 diabetes (T1D) when transferred into NOD-SCID mice^30,31^. For these experiments, we generated a specific CRM^pMHCII^ built with the mimotope peptide RLGL-WE14 in I-A^g7^ that binds the BDC2.5 TCR, and a negative control CRM^pMHCII^ built with a self-peptide from glucose phosphoisomerase (GPI282-294) presented in I-A^g7^ that is not associated with T1D^32,33^.

CD8^+^ T cells from NOD mice were activated *in vitro* and retrovirally transduced to express the ^5M^CARs. Phenotypic analysis showed that the ^5M^CAR-CTLs expressed the CRM^pMHCII^, as detected with anti-I-A^g7^ antibodies, and had an increase in CD80 levels compared with the natural levels of CD80 expression that are induced after activation (**sFig. 2a**)^34^. The majority of the ^5M^CAR-CTLs were CD44^hi^ CD62L^−^, Granzyme B^+^, Perforin^+^, FasL^lo^ and Fas^−^ (**Fig. S2b-d**), indicating they were primed CTLs with the potential to kill both by releasing cytotoxic granule proteins and by engaging Fas^35^.

To evaluate if these ^5M^CAR-CTLs could kill their targets in a TCR-specific manner we cultured specific (RLGL-WE14:I-A^g7^) and control (GPI:I-A^g7^) ^5M^CAR-CTLs with BDC2.5 CD4^+^ T cells at varying ratios of ^5M^CAR-CTL:target, and then enumerated the remaining targets after 16 hours culture. Here again we observed robust killing of the target CD4^+^ T cells by the specific ^5M^CAR-CTLs as compared to the control ^5M^CAR-CTLs. In contrast, neither ^5M^CAR-CTL population killed polyclonal CD4^+^ T cells from a NOD mouse in an off-target fashion (**Fig. 2a**). These results provide further evidence, with a second CRM^pMHCII^-TCR recognition model, that ^5M^CARs can redirect CTLs to specifically kill CD4^+^ T cells via recognition of the target T cell’s clonotypic TCR.

**Fig. 2:**
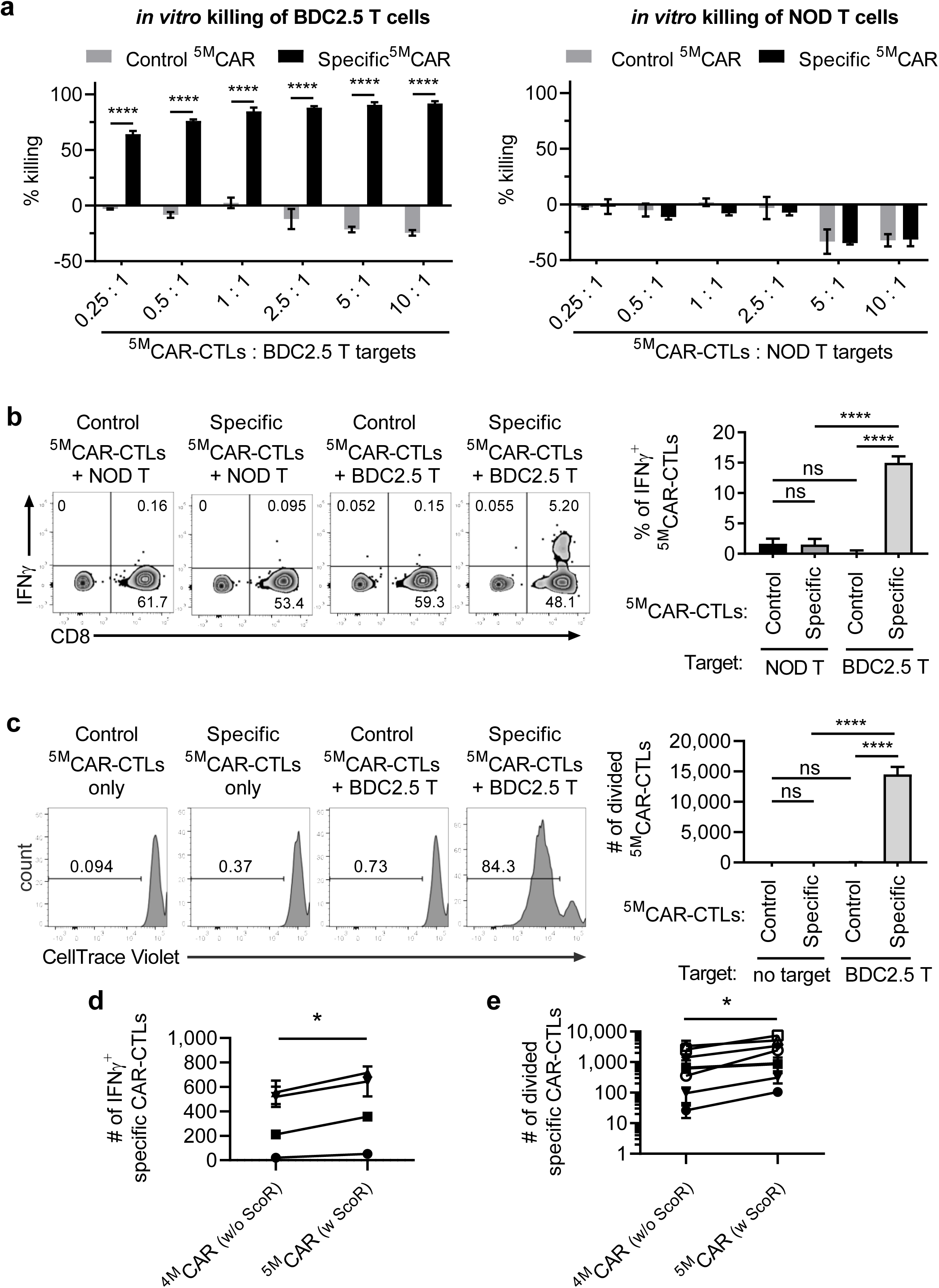
^5M^CARs redirect CTL function to target autoimmune CD4^+^ T cells. **a**, ^5M^CAR-CTL killing of target BDC2.5 CD4^+^ T cells (left) or control NOD CD4^+^ T cells (right) after co-culture with control (GPI:I-Ag7) or specific (RLGL-WE14:I-A^g7^) ^5M^CAR-CTLs is presented as in Fig. 1 (****p < 0.0001 by unpaired, two-tailed *t*-test)**. b**, IFN*γ* production by ^5M^CAR-CTLs. ^5M^CAR-CTLs incubated with target BDC2.5 T cells or NOD T cells for 6 hrs were stained for intracellular IFN*γ* and gated on live cells. Representative plots are shown. The bar graph shows the frequency of IFN*γ* producing ^5M^CAR-CTLs gated on GFP^+^CD8^+^mRasp^−^CD4^−^ cells (****p < 0.0001 by one-way ANOVA and Tukey’s post test). **c**, Proliferation of CellTrace violet labeled ^5M^CAR-CTLs after co-culture with or without target BDC2.5 T cells for 3 days. Representative histograms show CellTrace violet dilution of ^5M^CAR-CTLs. Bar graphs show the number of dividing ^5M^CAR-CTLs (****p < 0.0001 by one-way ANOVA and Tukey’s multiple comparison test). **d**, The number of specific ^4M^CAR-(without ScoR) or ^5M^CAR-CTLs (with ScoR) making IFN*γ* were measured by flow cytometry after coculture with target BDC2.5 T cells for 6 hrs. **e**, The number of divided specific ^4M^CAR-(without ScoR) or ^5M^CAR-CTLs (with ScoR) were by determined by flow cytometry after coculture with target BDC2.5 T cells for 3 or 4 days. Columns in **a-c** are show mean ± SD of triplicates or quadruplicates. Each graph is representative of 3 independent experiments. Lined data points in **d**,**e**, are shown as mean ± SD of triplicates or quadruplicates (*p < 0.05 by paired, two-tailed *t*-test). Each line represents an independent experiment.

Because CTL killing is reported to require less signaling than cytokine production or proliferation, we next asked if ^5M^CAR-CTLs make interferon γ (IFN γ) or proliferate in response to target CD4^+^ T cells^36^. Accordingly, we used intracellular cytokine staining and flow cytometry analysis to link IFNγ production to either the ^5M^CAR-CTLs or target T cells. After culturing specific and control ^5M^CAR-CTLs with either BDC2.5 CD4^+^ T cells or polyclonal CD4^+^ T cells from NOD mice for 6 hours we found that the specific ^5M^CAR-CTLs produced IFNγ only in response to the target BDC2.5 T cells, but not in response to the polyclonal NOD T cells, while the control ^5M^CAR-CTLs produced negligible IFNγ upon incubation with either BDC2.5 or NOD T cells (**Fig 2b**). We next tested whether ^5M^CAR-CTLs proliferate upon incubation with their targets. CellTrace Violet^TM^ (CTV)-labeled ^5M^CAR-CTLs were incubated for 3 days with either BDC2.5 or polyclonal NOD CD4^+^ T cells. Proliferating cells that diluted dye were enumerated by flow cytometry. Specific ^5M^CAR-CTLs only proliferated in response to BDC2.5 T cells, while the control ^5M^CAR-CTLs did not proliferate when incubated with either BDC2.5 or NOD T cells (**Fig 2c**). These data demonstrate that ^5M^CAR-CTLs can make cytokines and proliferate in response to CD4^+^ T cell targets expressing the appropriate TCR.

To complete our initial evaluation of ^5M^CAR performance, we tested the contribution of the ScoR module to ^5M^CAR-CTL function. The ScoR did not enhance killing of CD4^+^ T cell targets (**sFig. 2e**). Because killing is a lower-order function, requiring few cognate ligands (in this experiment there is a high ligand density of TCRs on the target cells), we next asked if the ScoR module impacted cytokine production or proliferation^11,36^. The ScoR did increase the number of specific ^5M^CAR-CTLs making IFNγ after 6 hours stimulation (**Fig. 2d**) and the number of specific ^5M^CAR-CTLs that divided after 3-4 days in culture (**Fig. 2e**). These data indicate that the ScoR makes a significant contribution to responses that might be useful *in vivo*. We therefore moved forward with the full ^5M^CAR system for *in vivo* analysis.

### ^5M^CAR-CTLs kill pathogenic CD4^+^ T cells *in vivo*

We next assessed the ability of specific (RLGL-WE14:I-A^g7^) and control (GPI:I-A^g7^) ^5M^CAR-CTLs to kill BDC2.5 CD4^+^ T cell targets by adapting a standard *in vivo* killing assay^37^. ^5M^CAR-CTLs were transferred into NOD mice. Twelve hours later the recipients received specific mRaspberry (mRasp)^+^ BDC2.5 CD4^+^ T cell targets mixed with control CTV-labeled NOD CD4^+^ T cell targets as a reference population. A separate cohort of NOD mice only received the mixture of targets (no ^5M^CAR-CTLs). The spleens were harvested 5.5 hours after target transfer and analyzed by flow cytometry (**sFig. 3**). Killing was evaluated by measuring changes in the frequency of specific targets relative to the reference targets (**Fig. 3**). The mice that did not receive ^5M^CAR-CTLs, and those that received control ^5M^CAR-CTLs had a mean ratio of 0.48 and 0.55 BDC2.5 to NOD CD4^+^ T cell targets, respectively, while those receiving specific ^5M^CAR-CTLs had a mean ratio of ~0.27, indicating ~50% killing of the target BDC2.5 CD4^+^ T cells relative to the control NOD CD4^+^ T cells (**Fig. 3**). These data show that ^5M^CAR-CTLs can rapidly find and eliminate their targets *in vivo*.

**Fig. 3:**
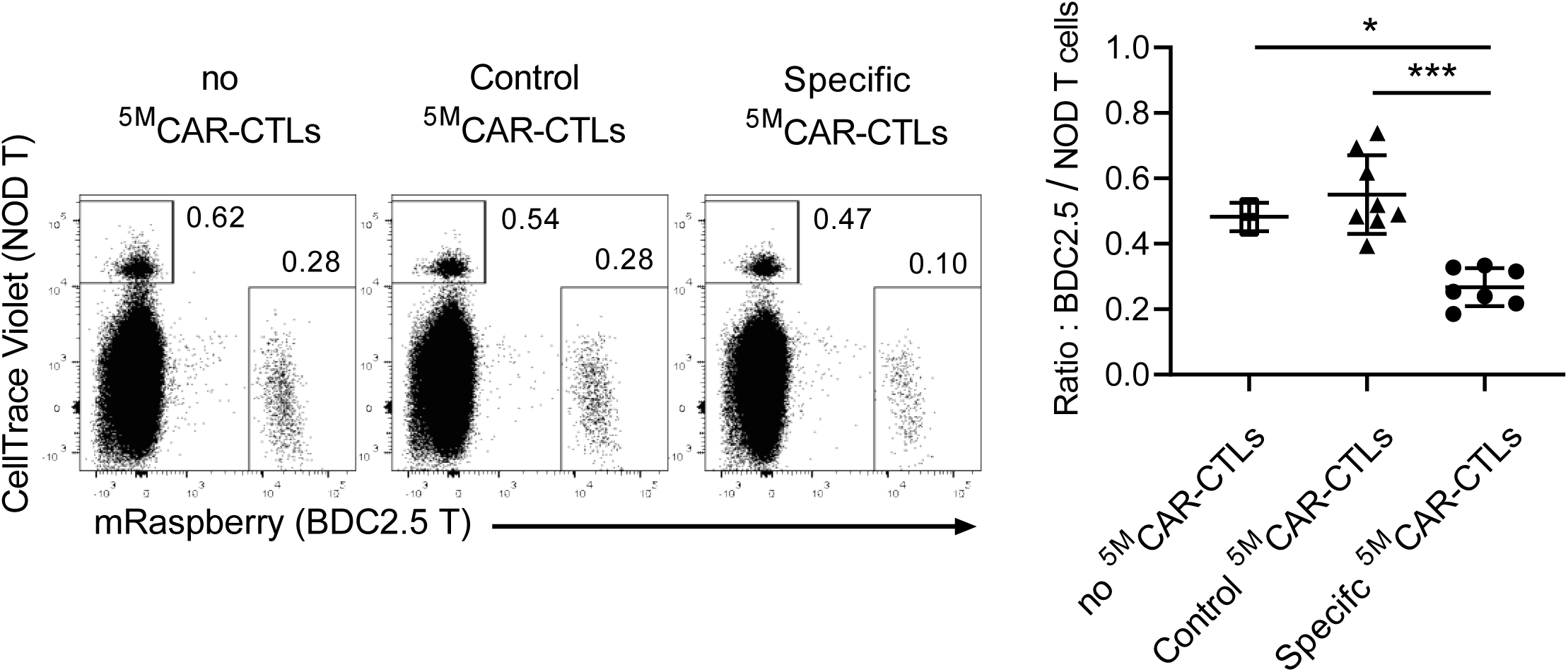
^5M^CAR-CTLs kill target T cells *in vivo*. Control (GPI:I-A^g7^) or specific (RLGL-WE14:I-A^g7^) ^5M^CAR-CTLs were adoptively transferred into NOD mice followed 12 hrs later with a mixture of mRasp^+^ BDC2.5 CD4^+^ T cells targets and CellTrace Violet-labeled NOD CD4^+^ T cells as a reference population. 5.5 hrs later the spleens were analyzed by flow cytometry to determine the extent of target cell killing. Plots show analysis of target and control cells in representative mice, gated live CD3^+^CD4^+^ cells (full gating shown in **Fig S3**). The graph shows the ratio of target cells/control cells as mean ± SD. Each point represents the ratio from a single spleen (*p < 0.05, ***p < 0.001 by one-way ANOVA and Tukey’s posttest). Data are shown as mean ± SD of combined from two independent experiments.

### ^5M^CAR-CTLs prevent autoimmune diabetes in NOD-SCID mice

The ability of ^5M^CAR-CTLs to kill autoimmune CD4^+^ T cells *in vivo* led us to ask if they could prevent BDC2.5 CD4^+^ T cell-mediated β-cell destruction and diabetes. For these experiments, we used a model in which we transferred BDC2.5 CD4^+^ T cells into NOD-SCID mice to induce diabetes^31^. In this model we have observed lymphocytic infiltration of the pancreas 5 days after BDC2.5 T cell transfer, which progresses to severe insulitis of most islets by day 6-8, deterioration of islet integrity, and diabetes onset between days 9-14 (**sFig. 4a,b**).

To explore if ^5M^CAR-CTLs can prevent diabetes we transferred BDC2.5 CD4^+^ T cells into NOD-SCID mice on day 0 (**Fig. 4a**). On day 1 the mice were divided into three cohorts: BDC2.5-only (untreated), control ^5M^CAR-CTL-treated, and specific ^5M^CAR-CTL-treated. Urine glucose was then measured daily to monitor diabetes onset and progression. All untreated and control ^5M^CAR-CTL-treated mice developed diabetes within 10 days, while all mice treated with specific ^5M^CAR-CTLs remained diabetes free for the duration of the 3-week experiment (**Fig. 4b**). Histological sections of the pancreases of the untreated and control ^5M^CAR-CTL-treated diabetic mice showed widespread lymphocytic infiltration and islet destruction, while pancreases from mice treated with specific ^5M^CAR-CTLs appeared free of infiltration (**Fig. 4c**). Disease amelioration corresponded with nearly complete elimination of BDC2.5 CD4^+^ T cells in the spleens of the specific ^5M^CAR-CTL-treated mice, as evaluated by flow cytometry (**Fig. 4d,e**). Importantly, the ^5M^CAR-CTLs were present in the spleens of the recipient mice 21 days after transfer, indicating that specific ^5M^CAR-CTLs have the potential for long-term engraftment (**Fig. 4d,e**).

**Fig. 4:**
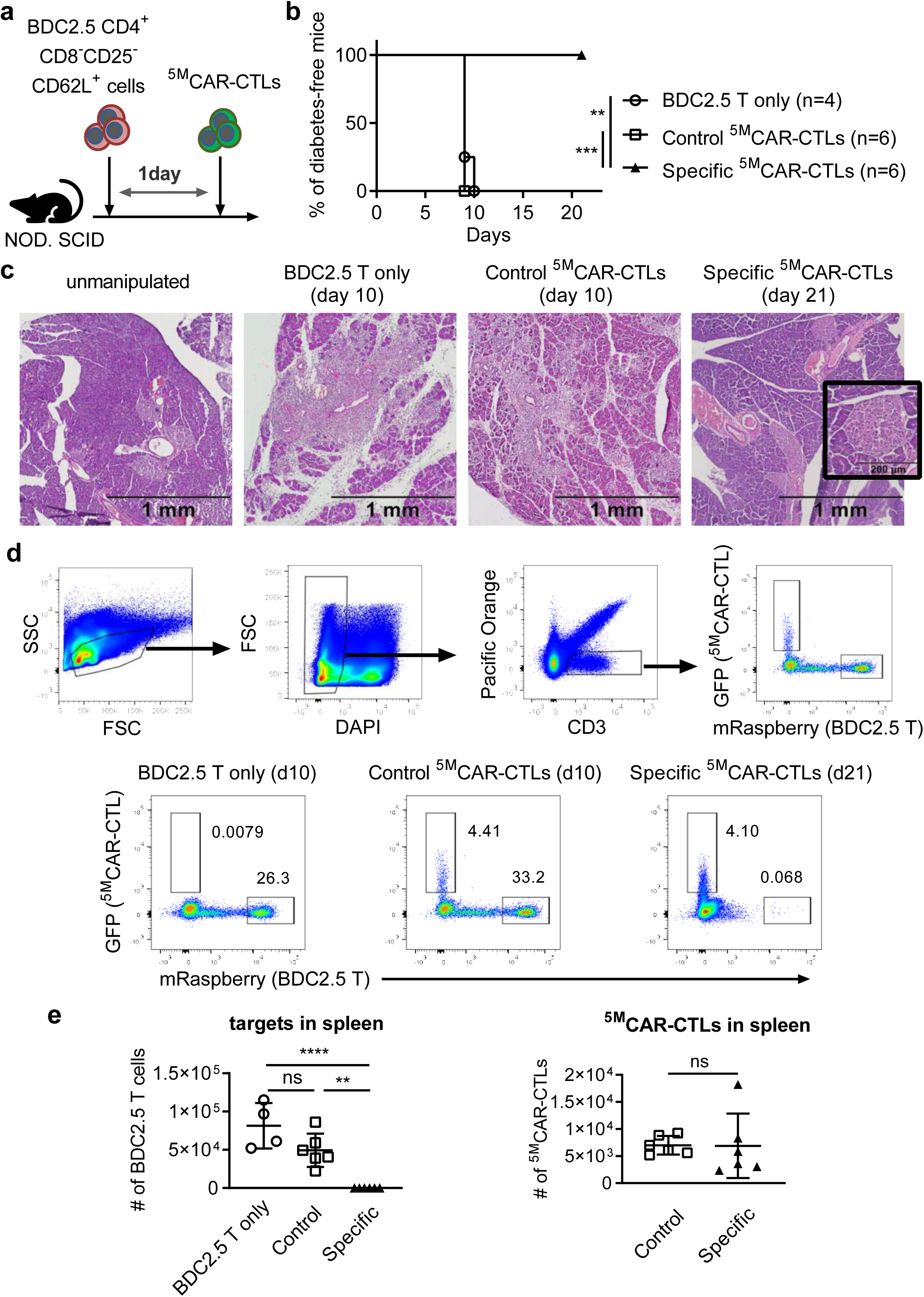
^5M^CAR-CTLs prevent BDC2.5 CD4^+^ T cell-induced T1D in NOD-SCID mice. **a**, BDC2.5 CD4^+^ T cells were adoptively transferred into NOD-SCID mice on day 0. On day 1 the mice were either treated with ^5M^CAR-CTLs or left untreated. **b**, Survival curve shows the percentage of diabetes-free mice that were treated with either control (GPI:I-A^g7^) ^5M^CAR-CTLs, specific (RLGL-WE14:I-A^g7^) ^5M^CAR-CTLs, or left untreated (BDC2.5 only) (**p < 0.01, ***p < 0.001 by log-rank test). **c**, Pancreases of representative mice from each group are shown stained with hematoxylin-eosin (Magnification: 4X). Black box inset shows clear islet (20X, scale bar = 200μm). **d**,**e**, Analysis of mRasp^+^ target CD4^+^ T cells and GFP^+ 5M^CAR-CTLs in spleens of treated and untreated mice. **d**, Gating schematic for the analysis. **e**, Representative dot plots show frequencies of targets and ^5M^CAR-CTLs in spleens. Graphs show absolute cell counts of BDC2.5 CD4^+^ T cells or ^5M^CAR-CTLs. Each point represents an individual recipient. Horizontal lines indicate mean ± SD [**p < 0.01, ****p < 0.0001 by one-way ANOVA and Tukey’s posttest (bottom, left) or unpaired, two-tailed *t*-test (bottom, right). ns means not statistically significant. Data are representative of two similar independent experiments. Numbers of mice/group are indicated in the figure.

### ^5M^CAR-CTLs halt ongoing insulitis

Having established that BDC2.5-specific ^5M^CAR-CTLs can eliminate their targets and prevent diabetes in NOD-SCID mice when transferred one day after BDC2.5 CD4^+^ T cells, we asked if these ^5M^CAR-CTLs could reverse insulitis and prevent diabetes if transferred at a later stage in disease progression. Specifically, we asked whether ^5M^CAR-CTLs could prevent diabetes on day 7 after BDC2.5 transfer, at which point islets are invaded by lymphocytic infiltrates and they have substantially lost their structural integrity (**Fig. 5a, sFig 4a**).

**Fig. 5:**
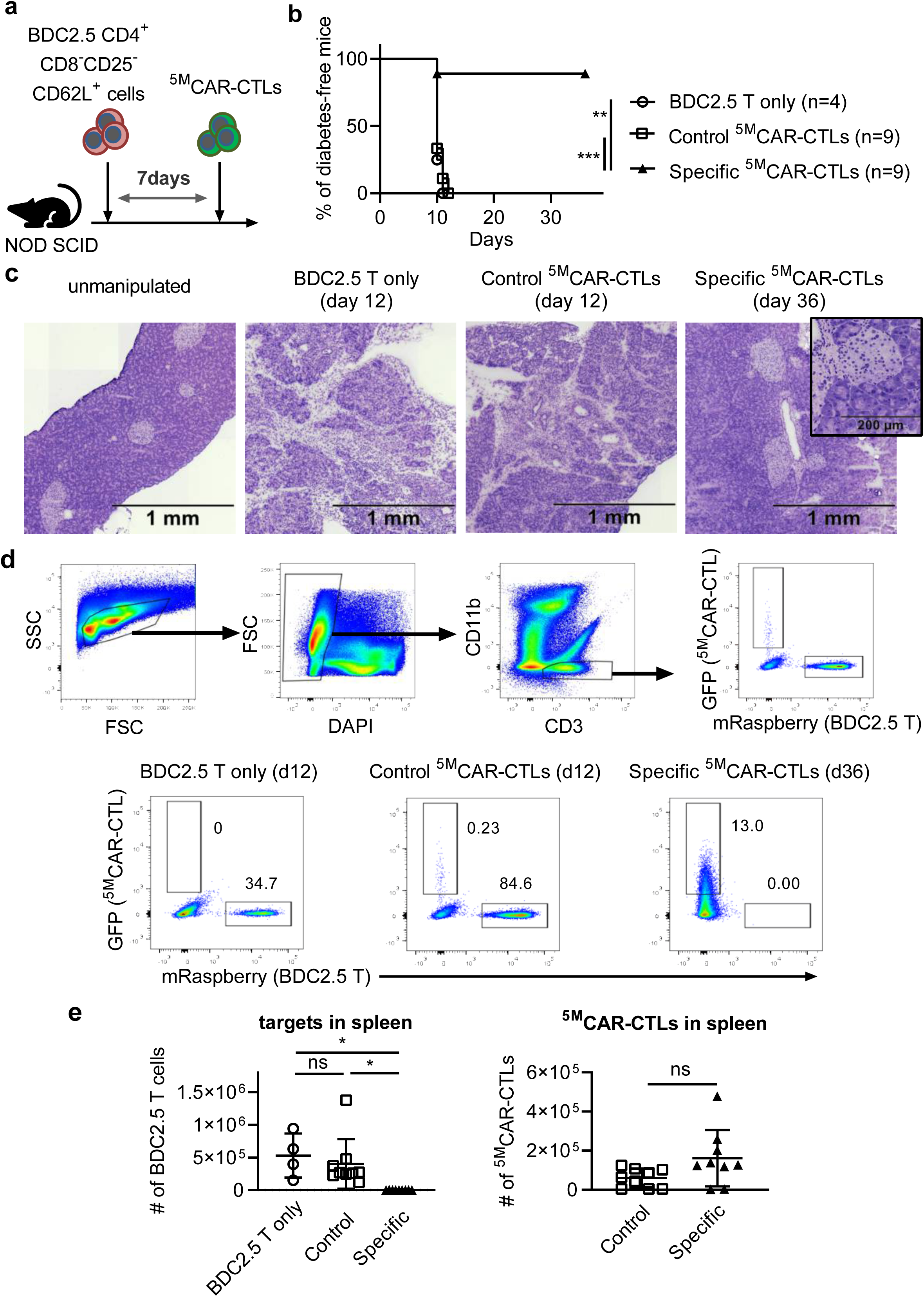
^5M^CAR-CTLs prevent diabetes after initiation of insulitis. NOD-SCID mice receiving BDC2.5 CD4^+^ T cells on day 0 were treated with ^5M^CAR-CTLs on day 7, or left untreated. **b**, Survival curve shows the percentage of diabetes-free mice treated with either control (GPI:I-A^g7^) ^5M^CAR-CTLs, specific (RLGL-WE14:I-A^g7^) ^5M^CAR-CTLs, or left untreated (BDC2.5 only) (**p < 0.01, ***p < 0.001 by log-rank test). **c**, Pancreases of representative mice from each group, stained with hematoxylin-eosin, are shown (Magnification: 4X). Black box inset shows clear islet (20X, scale bar = 200μm). **d**,**e**, Analysis of mRasp^+^ CD4^+^ T cells and GFP^+ 5M^CAR-CTLs in spleens of treated and untreated mice. **d**, Gating schematic for the analysis. **e**, Representative dot plots show frequencies of targets and ^5M^CAR-CTLs in spleens. Graphs show absolute cell counts of BDC2.5 T cells or ^5M^CAR-CTLs. Each data point represents an individual recipient. Horizontal lines indicate mean ± SD. *p < 0.05 by one-way ANOVA and Tukey’s multiple comparison test (bottom, left) or by unpaired, two-tailed *t*-test (bottom, right). ns means not statistically significant. Data are combined from two independent experiments. Numbers of mice/group are indicated in the figure.

BDC2.5 T cells were transferred into NOD-SCID mice on day 0 and the recipients were divided into three cohorts on day 7: BDC2.5-only (untreated), control ^5M^CAR-CTL-treated, and specific ^5M^CAR-CTL-treated. All untreated and control ^5M^CAR-CTL-treated mice developed diabetes by day 13, while 8 of 9 specific ^5M^CAR-CTL-treated mice remained diabetes free (**Fig. 5b**). Those protected from diabetes maintained normal weight and blood glucose throughout the experiment (**sFig 5a,b**). Furthermore, histological sections from the pancreases of specific ^5M^CAR-CTL-treated mice showed a lack of insulitis, whereas the structural integrity of the islets was destroyed in the untreated and control ^5M^CAR-CTL-treated mice (**Fig 5c**). Importantly, at the experimental endpoints (day 11-13 for mice with diabetes and day 36 for diabetes free mice) analysis of the spleens and pancreatic lymph nodes (pLN) indicated that the BDC2.5 CD4^+^ T cells had been eliminated (**Fig. 5d,e and sFig. 5c**). Even in the one specific ^5M^CAR-CTL-treated animal that had become diabetic, the BDC2.5 CD4^+^ T cells had been nearly eliminated by day 11; although apparently this was too late for diabetes protection in this recipient (**sFig. 5c**). ^5M^CAR-CTLs also remained in the spleens and pLN of recipient mice for the duration of the experiment (**Fig. 5e, sFig. 5c**), again demonstrating that they can engraft for several weeks.

### ^5M^CAR-CTLs home to and persist in the pancreas

Having determined that specific ^5M^CAR-CTLs can reverse insulitis and prevent diabetes, while control ^5M^CAR-CTLs cannot, we evaluated where the ^5M^CAR-CTLs traffic, where they encounter their targets, and how the numbers of both the targets and ^5M^CAR-CTLs change over time. Here again NOD-SCID mice received BDC2.5 CD4^+^ T cells on day 0 and ^5M^CAR-CTLs on day 7. Cohorts of mice were euthanized on day 7 (pre-^5M^CAR-CTL transfer), 8, 10, 15, and 36 to enumerate BDC2.5 CD4^+^ T cells and ^5M^CAR-CTLs within the spleens, pLNs, and pancreases.

In mice treated with control ^5M^CAR-CTLs, the number of BDC2.5 CD4^+^ T cells in the spleens and pLN remained relatively constant from day 7-10, but increased from day 10 to 15 during the time in which the mice developed diabetes (**Fig. 6a and sFig. 6**). In the pancreas, BDC2.5 CD4^+^ T cells increased in numbers by day 10 and, by day 15, had expanded ~100-fold compared to day 7 (**Fig. 6b**).

**Fig. 6:**
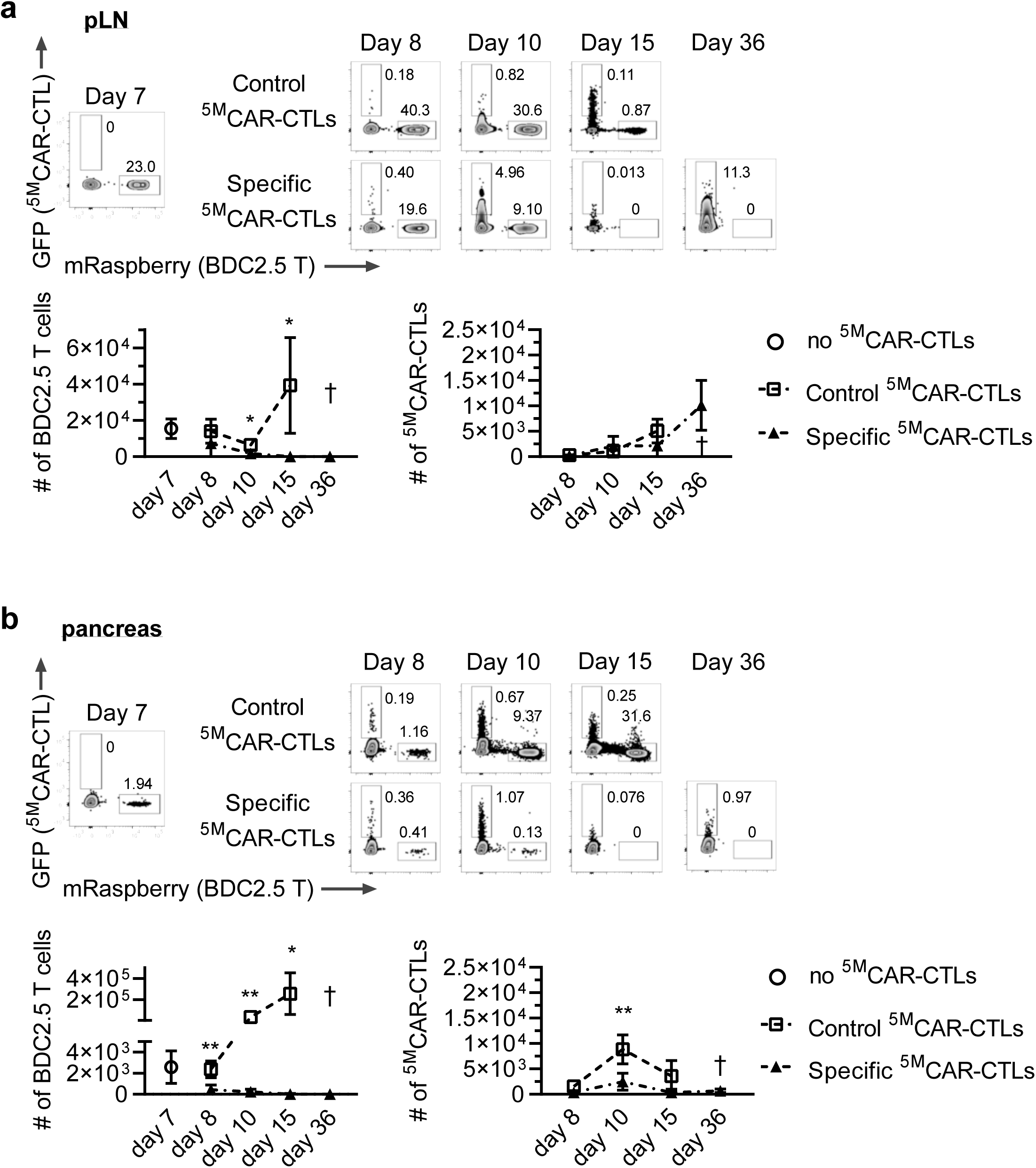
^5M^CAR-CTLs migrate to the pancreas and eliminate BDC2.5 CD4^+^ T cells. NOD-SCID mice receiving BDC2.5 T cells (day 0) were either treated with control (GPI:I-A^g7^) ^5M^CAR-CTLs, specific (RLGL-WE14:I-A^g7^) ^5M^CAR-CTLs, or euthanized prior to treatment on day 7. Treated groups were euthanized on day 8, 10, 15 or 36. Flow cytometry was used to determine the frequency and number of mRasp^+^ CD4^+^ BDC2.5 T cells and GFP^+ 5M^CAR-CTLs in the pLNs (**a**) or pancreases (**b**). **a**.**b**, Representative dot plots show frequencies of target BDC2.5 T cells, and ^5M^CAR-CTLs (top panels, pre-gated on live, CD11b^−^ cells). Graphs show the number of CD4^+^ BDC2.5 T cells (left) or change of the number of ^5M^CAR-CTLs (right). Data show combined results from two independent experiments as mean ± SD. 3-6 mice were analyzed for each group and time point (*p < 0.05, **p < 0.01 by unpaired, two-tailed *t*-test between the control and specific ^5M^CAR-CTLs groups). † means no data. Numbers of mice/group are indicated in the figure.

In mice treated with specific ^5M^CAR-CTLs, a reduction in the frequency and number of BDC2.5 CD4^+^ T cells was evident by day 10 in the pLNs, in the spleens by day 15, and as early as day 8 in the pancreases. Importantly, BDC2.5 CD4^+^ T cells were barely detectable by day 15 in any of the analyzed tissues, and none were detected by the experimental endpoint at day 36.

Regarding the ^5M^CAR-CTLs, both the control and specific populations were detected in the spleen and the pLN on day 8 (**Fig. 6a and sFig. 6**). Importantly, both populations also homed to the inflamed pancreases within the first 24 hours post-transfer (day 8)(**Fig. 6b**). The numbers of control ^5M^CAR-CTLs stayed relatively constant in the spleens from days 8 to 15, increased slightly over time in the pLNs, and actually peaked at day 10 in the pancreas while the number of specific ^5M^CAR-CTLs remained relatively constant in the spleens, pLNs, and pancreases from day 8 to 15. Of note, the number of specific ^5M^CAR-CTLs was significantly lower than the control ^5M^CAR-CTLs at day 10 and 15 in the pancreases, mirroring the loss of BDC2.5 cells. Finally, the specific ^5M^CAR-CTLs were present in the spleens, pLNs, and pancreases at the termination of the experiment on day 36.

Overall, these results indicate that ^5M^CAR-CTLs can rapidly home to inflamed pancreases, as well as to lymphoid tissues, and can eliminate their target cells from those tissues in a TCR-specific fashion. Furthermore, they can persist in these tissues for weeks, offering the potential for long-term tissue-resident protection.

### A panel of ^5M^CAR-CTLs can prevent autoimmune diabetes in NOD mice

Once we established that ^5M^CAR-CTLs can prevent T1D induction by a monoclonal population of pathogenic CD4^+^ T cells in NOD-SCID mice, we wanted to test the efficacy of ^5M^CAR-CTL treatment in the NOD mouse model of spontaneous T1D. In this model insulitis can be detected by 10 weeks of age in most mice, while diabetes onset typically occurs at 12-14 weeks of age^38,39^. There is a sexual dimorphism regarding the frequency of disease incidence, with 80% or more of female mice becoming diabetic by 45 weeks^38,40^. Also, in NOD mice as in humans, T1D is thought to be initiated by T cells reactive to one or a small number of self-pMHC^41,42^. For example, mice that lack a key amino acid in an insulin peptide (INSB9-23) presented by I-A^g7^ are protected from diabetes development, suggesting that this peptide is critical to disease initiation^42^. In addition, CD4^+^ T cells specific for newly discovered hybrid insulin peptides (HIP) presented by I-A^g7^, including the BDC2.5 T cells used here, can induce diabetes in adoptive transfer models^43^. We therefore generated three additional CRM^pMHCII^, presenting either the insulin peptide (INSB9-23) or two hybrid insulin peptides (HIP2.5, HIP6.9) in I-A^g7^, to test if a mixture of ^5M^CAR-CTLs could prevent diabetes in NOD mice. Of note, the HIP2.5:I-A^g7^- and RLGL-WE14:I-A^g7^-based CRM^pMHCII^ used above are likely to target partially overlapping cohorts of autoimmune CD4^+^ T cells^44^.

To determine if ^5M^CAR-CTLs that target CD4^+^ T cells specific to these self-pMHCII can establish long-term engraftment and protect against autoimmune diabetes, we transferred a mixture of specific (INSB:I-A^g7^, HIP2.5:I-A^g7^, HIP6.9:I-A^g7^ and RLGL-WE14:I-A^g7^) or control (GPI:I-A^g7^) ^5M^CAR-CTLs into neonatal male and female NOD mice. We chose neonatal mice as recipients because: 1) disease-initiating autoimmune CD4^+^ T cells are thought to emerge from the thymus soon after birth; 2) physiological β-cell death at 2 weeks of age is reported to trigger priming of self-reactive T cells; and, 3) we wanted to eliminate the target CD4^+^ T cells before they could provide help to autoimmune CD8^+^ T cells^45,46^. A third group of newborn mice was left untreated to establish a baseline for comparison of T1D incidence.

We first asked if ^5M^CAR-CTLs would persist throughout the critical period of early autoimmune T cell development, early insulitis, and β cell apoptosis. To address this question, we sacrificed a cohort of male mice at 13 weeks of age to assess ^5M^CAR-CTL engraftment. ^5M^CAR-CTLs were detected in both the control and specific groups of mice (**Fig. 7a**). These data verified that the ^5M^CAR-CTLs can engraft long-term in NOD mice and were thus present during the period of life that is thought to be critical for spontaneous T1D in these animals.

**Fig. 7:**
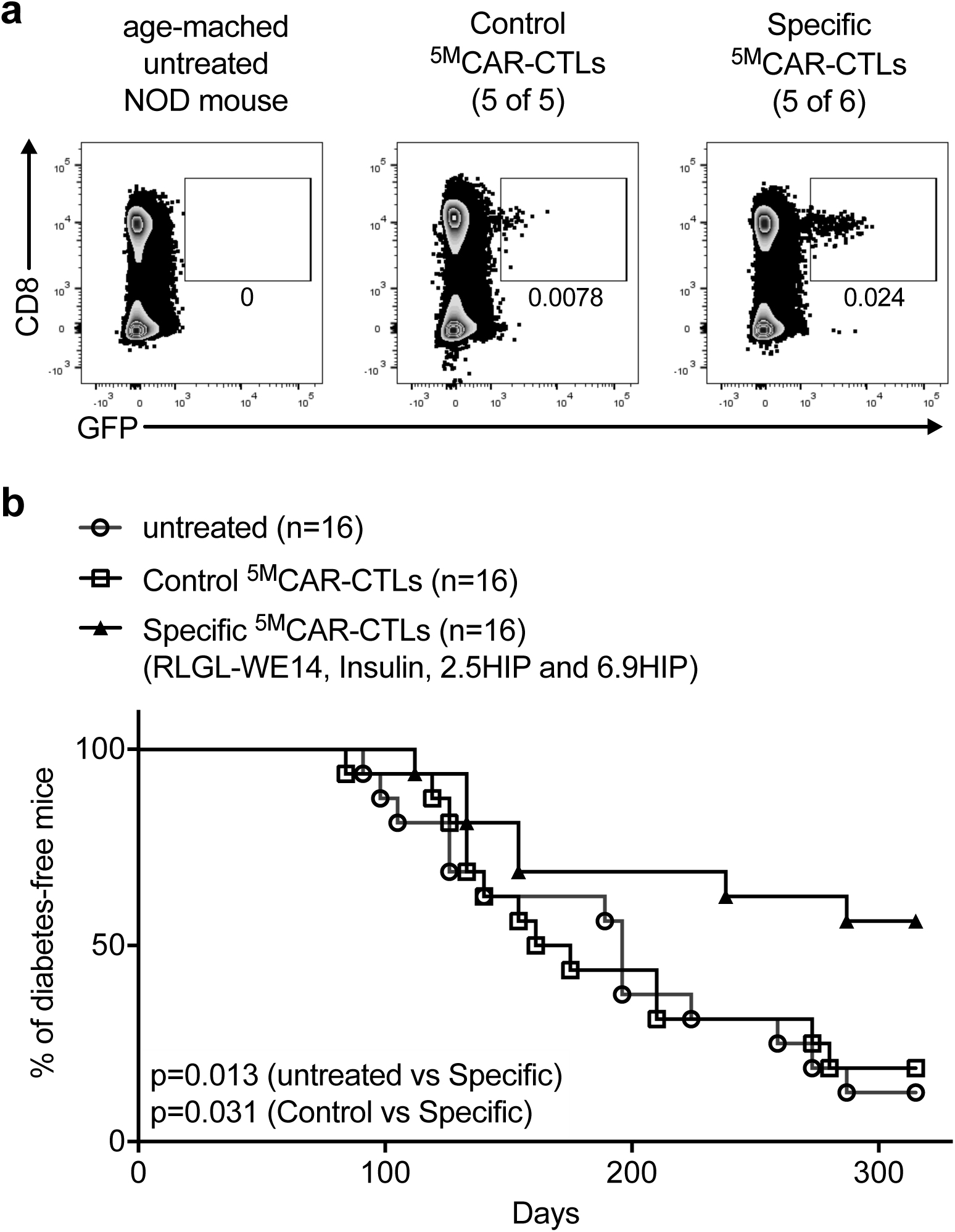
Treatment with an oligoclonal set of specific ^5M^CAR-CTLs decreases diabetes incidence in NOD mice. Newborn NOD mice received control (GPI:I-A^g7^) ^5M^CAR-CTLs, a mixture of specific ^5M^CAR-CTLs targeting four populations of T1D-related autoimmune T cell (INSB:I-A^g7^, HIP2.5:I-A^g7^, HIP6.9:I-A^g7^ and RLGL-WE14:I-A^g7^), or were left untreated. **a**, A subset of male mice were euthanized at 13 weeks of age, and their spleens were analyzed by flow cytometry. ^5M^CAR-CTLs were identified by CD8 and GFP expression (pre-gated on live, CD3^+^ cells). Plots from representative engrafted mice are shown (5/5 control ^5M^CAR-CTL recipients engrafted, 5/6 specific ^5M^CAR-CTL recipients engrafted). **b**, Female mice were screened weekly for glycosuria from day 32 to day 315 and cumulative incidence of diabetes (survival probability plots based on the Kaplan-Meier method) are shown with exact p values determined by log-rank test. Data show the combined results from three independent cohorts. Numbers of mice/group are indicated in the figure.

Due to the sexual dimorphism of T1D in NOD mice, females were followed for 315 days to assess diabetes development. Diabetes onset and progression for those treated with control ^5M^CAR-CTLs mirrored that of the untreated group in the Kaplan-Meier plots (**Fig. 7b**). At day 315, 14 of 16 untreated mice had developed diabetes and 13 of 16 control ^5M^CAR-CTL-treated mice had developed diabetes. In contrast, fewer mice treated with the specific ^5M^CAR-CTLs developed diabetes (7 of 16) when compared to the untreated animals or those that were treated with control ^5M^CAR-CTLs. These data demonstrate that a mixture of ^5M^CAR-CTLs targeting autoimmune CD4^+^ T cells with a limited specificity can significantly reduce the incidence of T1D in NOD mice.

We also analyzed the data using the Cox proportional hazards model. Risk of diabetes incidence in mice with the specific ^5M^CAR-CTLs were approximately 3 times lower when compared with the untreated and control mice. Hazard ratios (HRs) and the respective confidence intervals (CI) for the specific group in comparison to the untreated group were HR (95% CI): 0.33 (0.13, 0.82); p value=0.017; and the specific group in comparison to the control group were HR (95% CI): 0.36 (0.14, 0.91); p value=0.030.

Finally, there were no differences in the diabetes incidence between the control and untreated groups HR (95% CI): 0.91 (0.43, 1.94); p value = 0.808. Finally, a cohort of surviving mice (4 males and 5 females) were euthanized after day 315 for analysis of pooled spleens and lymph nodes. We detected ^5M^CAR-CTLs in 2 out of 9 mice (**sFig. 7a**). To explore if additional ^5M^CAR-CTLs might be present below the limit of detection we transferred BDC2.5 CD4^+^ T cells into the remaining diabetes-free mice to potentially expand out the BDC2.5-specific ^5M^CAR-CTLs. In these boosted mice, 2 of 4 specific ^5M^CAR-CTL recipient mice had detectable specific ^5M^CAR-CTLs (**sFig. 7b**). We cannot know if the ^5M^CAR-CTLs in the specific group expanded in response to the BDC2.5 T cells, or would have been detectable without them. Nevertheless, the data indicate that ^5M^CAR-CTLs engraft up to a year or longer in a subset of animals despite the CD8^+^ T cell attrition that normally occurs with aging^47^.

## Discussion

The goals of this study were to evaluate the function of our 1^st^ generation biomimetic ^5M^CAR *in vitro* and *in vivo*. We show that ^5M^CARs can redirect CTL specificity against clonotypic TCRs expressed by CD4^+^ T cells and can co-opt CTL functions, including IFNγ production, proliferation, and killing, in a TCR-specific manner *in vivo*. We also demonstrate that ^5M^CAR-CTLs can rapidly eliminate pathogenic CD4^+^ T cells, and thus neutralize their detrimental impact, in mouse models of T1D. The ^5M^CAR design, biological implications of the data, and potential applications of this technology are discussed below along with some thoughts on the future of the technology.

Our biomimetic ^5M^CAR design integrates three key operating principles not found together in other CARs. First, the receptor modules assemble with the CD3 signaling modules into complexes that possess a full complement of 10 ITAMs, as well as other key motifs described elsewhere, that are thought to either mediate or regulate signaling through the TCR-CD3 complex^12,21,48–52^. This feature was incorporated into the design because the multiplicity of ITAMs that get phosphorylated during signaling via the TCR, or even ^1M^CARs, influences subsequent T cell function; furthermore, we speculate that preserving access to the natural signaling apparatus may serve as an important safety feature to prevent dysregulated responses, as improper signaling can lead to dysregulated T cell function^21,53–55^. We note that a recently reported 4-module TruC CAR also takes advantage of the native CD3 signaling modules^56^. For the second key design principle we maintained the natural biophysical properties of receptor:ligand interactions that have evolved to mediate T cell activation. T cells are normally selected and activated within a narrow kinetic window of TCR-pMHC interactions, while higher than normal affinity leads to attenuated or even undesired effects; therefore, tuning the affinity of a CAR for its ligand to this natural range, while maintaining specificity, is an important consideration for optimizing function^19,57–60^. Of note, a 4-module approach was previously described whereby the variable regions of the anti-TNP SP6 antibody were used to make a chimeric TCR (cTCR) that associated with the CD3 signaling modules, and redirected CTL specificity and function *in vitro*^61^. In that report the SP6:TNP interaction that was used was of very low affinity^62^, and thus more akin to TCR:pMHCII interactions (µM) than typical antibody:antigen interactions (nM), suggesting affinity optimized receptor modules based on antibodies might be effective in a next generation ^5M^CAR approach. Finally, our biomimetic design is unique from other CAR designs in that it includes a surrogate coreceptor module. CD4 and CD8 sequester Lck away from the CD3 ITAMs in the absence of cognate pMHC, meaning that they should naturally keep Lck away from ITAMs associated with CARs of any design and thus prevent optimal CAR signaling^17^. Our ScoR provides the opportunity for Lck to be positioned proximally to the CRM^pMHCII^-CD3 ITAMs within the immunological synapse between a ^5M^CAR-CTL and a CD28^+^ or CTLA-4^+^ target T cell. An appealing conceptual aspect of using a ScoR is that alternate designs could be used to enhance targeting if needed. For example, ScoRs could be built with the ectodomains of other molecules that localize to the center of the immunological synapse (e.g. PD-L1 or CD70) to target the subset of CD8^+^ CD28^−^ Tm cells that accumulate in humans with aging^63^. Altogether, the data presented here demonstrate that the three operating principles discussed above can be successfully integrated into a functional ^5M^CAR system.

While the current study focused on engineering and testing our ^5M^CARs, we also made some noteworthy observations that speak to the usefulness of ^5M^CAR-CTLs as tools for studying the role of particular T cell populations in biology. For example, we show that ^5M^CAR-CTLs can rapidly home to the site of inflammation, eliminate their target population in a TCR-specific manner, and persist for months in mice. We also show that treating neonatal NOD mice with a mixture of ^5M^CAR-CTLs that target a limited number of known pMHCII-specific autoimmune CD4^+^ T cells can significantly decrease diabetes incidence, consistent with the idea that one or a limited number self-pMHCII reactivities initiate disease. Together, these data indicate that ^5M^CAR-CTLs can be used to eliminate specific T cell populations in mice and then study how particular immune responses are initiated and proceed in their absence.

In addition to their potential research applications, one benefit of using mouse models of T1D as testing grounds for evaluating ^5M^CAR-CTL function is that our results point to therapeutic applications for treating diseases mediated by pathogenic T cells. For example, we show that ^5M^CAR-CTLs can reverse ongoing insulitis and dramatically decreases T1D disease incidence in NOD mice. A prior ^1M^CAR study showed that targeting a single pathogenic CD8^+^ T cell population can also decrease diabetes incidence in NOD mice^64^; therefore, targeting a mixture of CD4^+^ and CD8^+^ T cell clonotypes with ^5M^CAR-CTLs in patients with preclinical disease (e.g. two autoantibodies), who are likely to develop diabetes, may be effective as a preventative therapy^65^. Multiple sclerosis (MS) is another autoimmune disease involving pathogenic CD4^+^ T cells for which dominant MHCII associations exist and a number of self-peptide auto-antigens have been described^66^. Importantly, MS cannot be managed as readily as diabetes, the methods for management are immunosuppressive, and ^1M^CARs have shown efficacy in experimental autoimmune encephalitis (EAE), a mouse model of MS^67^. Given the success of ^1M^CAR-T cell therapy in treating hematologic malignancies, another prime target for ^5M^CAR-CTL therapy would be TCR^+^ leukemias and lymphomas of the T cell lineage (e.g. T-ALL and CTCL) which are typically clonal and thus would not require targeting multiple specificities^68,69^. Significant advances have been made in identifying pMHC that specifically interact with a TCR^70,71^. Using such methods to identify a pMHC that binds a T lymphoma-derived TCR would enable the rapid generation of ^5M^CAR-CTLs to target TCR^+^ tumors. Finally, ^5M^CAR-CTLs could potentially be used to protect transplanted tissue if the appropriate pathogenic T cell clonotypes that mediate rejection can be eliminated prior to transplant^72^. Importantly, the lack of weight loss or overt signs of distress by the mice used in our studies suggest that the ^5M^CAR-CTLs themselves are not pathogenic. In addition, the conservation of pMHC and TCR structures between mice and humans suggest that humanized ^5M^CARs should function similarly.

Moving forward, a multi-pronged approach is required to advance biomimetic ^5M^CAR designs. Directly comparing their performance with other CAR designs, as well as their natural counterparts, will provide benchmarks for the iterative process of refinement to enhance their function according to the roadmap that has advanced ^1M^CARs^7–9^. Additional basic research into the molecular machinery that naturally drives T cell activation is also of fundamental importance, as it will provide a more complete blueprint for the development of future generations of biomimetic designs. These lines of investigation will provide us with the information needed to refine our 1^st^ generation biomimetic ^5M^CARs, optimize their application, and design future generations of ^5M^CAR modules that are tailored to target specific cell populations.

## Methods

### ^5M^CAR construction

^5M^CAR constructs were generated by standard molecular biology techniques. The genes encoding the CRM^pMHCII^ and ScoR were cloned into pUC18 (Fermentas), sequenced (ELIM BIOPHARM), and then subcloned into an MSCV-based retroviral expression vector.

The following sequence for the chimeric CD80-Lck surrogate coreceptor (ScoR) was subcloned into the “pP2” puromycin-resistance MSCV vector (MCS-IRES-Puro resistance^73^) via 5’XhoI and 3’EcoRI:

> acgtctagatacctcgaggccaccatggcttgcaattgtcagttgatgcaggatacaccactcctcaagtttccatgtccaaggctcattcttctctttgtgctgctgattcgtctttcacaagtgtcttcagatgttgatgaacaactgtccaagtcagtgaaagataaggtattgctgccttgccgttacaactctcctcatgaagatgagtctgaagaccgaatctactggcaaaaacatgacaaagtggtgctgtctgtcattgctgggaaactaaaagtgtggcccgagtataagaaccggactttatatgacaacactacctactctcttatcatcctgggcctggtcctttcagaccggggcacatacagctgtgtcgttcaaaagaaggaaagaggaacgtatgaagttaaacacttggctttagtaaagttgtccatcaaagctgacttctctacccccaacataactgagtctggaaacccatctgcagacactaaaaggattacctgctttgcttccgggggtttcccaaagcctcgcttctcttggttggaaaatggaagagaattacctggcatcaatacgacaatttcccaggatcctgaatctgaattgtacaccattagtagccaactagatttcaatacgactcgcaaccacaccattaagtgtctcattaaatatggagatgctcacgtgtcagaggacttcacctgggaaaaacccccagaagaccctcctgatagcaagaacacacttgtgctctttggggcaggattcggcgcagtaataacagtcgtcgtcatcgttgtcatcatcaaatgcttctgtaagcacagaagctgtttcagaagaaatgaggcaagcagagaaacaaacaacagccttaccttcgggcctgaagaagcattagctgaacagaccgtcttccttaccactagtcactatcccatagtcccactggacagcaagatctcgctgcccatccggaatggctctgaagtgcgggacccactggtcacctatgagggatctctcccaccagcatccccgctgcaagacaacctggttatcgccctgcacagttatgagccctcccatgatggagacttgggctttgagaagggtgaacagctccgaatcctggagcagagcggtgagtggtggaaggctcagtccctgacgactggccaagaaggcttcattcccttcaacttcgtggcgaaagcaaacagcctggagcctgaaccttggttcttcaagaatctgagccgtaaggacgccgagcggcagcttttggcgcccgggaacacgcatggatccttcctgatccgggaaagcgaaagcactgcggggtccttttccctgtcggtcagagacttcgaccagaaccagggagaagtggtgaaacattacaagatccgtaacctagacaacggtggcttctacatctcccctcgtatcacttttcccggattgcacgatctagtccgccattacaccaacgcctctgatgggctgtgcacaaagttgagccgtccttgccagacccagaagccccagaaaccatggtgggaggacgaatgggaagttcccagggaaacactgaagttggtggagcggctgggagctggccagttcggggaagtgtggatggggtactacaacggacacacgaaggtggcggtgaagagtctgaaacaagggagcatgtcccccgacgccttcctggctgaggctaacctcatgaagcagctgcagcacccgcggctagtccggctttatgcagtggtcacccaggaacccatctacatcatcacggaatacatggagaacgggagcctagtagattttctcaagactccctcgggcatcaagttgaatgtcaacaaacttttggacatggcagcccagattgcagagggcatggcgttcatcgaagaacagaattacatccatcgggacctgcgcgccgccaacatcctggtgtctgacacgctgagctgcaagattgcagactttggcctggcgcgcctcattgaggacaatgagtacacggcccgggagggggccaaatttcccattaagtggacagcaccagaagccattaactatgggaccttcaccatcaagtcagacgtgtggtccttcgggatcttgcttacagagatcgtcacccacggtcgaatcccttacccaggaatgaccaaccctgaagtcattcagaacctggagagaggctaccgcatggtgagacctgacaactgtccggaagagctgtaccacctcatgatgctgtgctggaaggagcgcccagaggaccggcccacgtttgactaccttcggagtgttctggatgacttcttcacagccacagagggccagtaccagccccagcctggtacctagtgagaattctacatg

The following sequence for the I-E^k^α.TCRα chimeric CRM^pMHCII^*^α^* subunit was subcloned into the “pZ4” zeocin-resistance MSCV vector (MCS-IRES-Zeo^73^) via 5’XhoI and 3’BamHI:

> aataagcttctcgagcgccaccatggccacaattggagccctgctgttaagatttttcttcattgctgttctgatgagctcccagaagtcatgggctatcaaagaggaacacaccatcatccaggcggagttctatcttttaccagacaaacgtggagagtttatgtttgactttgacggcgatgagattttccatgtagacattgaaaagtcagagaccatctggagacttgaagaatttgcaaagtttgccagctttgaggctcagggtgcactggctaatatagctgtggacaaagctaacctggatgtcatgaaagagcgttccaacaacactccagatgccaacgtggccccagaggtgactgtactctccagaagccctgtgaacctgggagagcccaacatcctcatctgtttcattgacaagttctcccctccagtggtcaatgtcacctggttccggaatggacggcctgtcaccgaaggcgtgtcagagacagtgtttctcccgagggacgatcacctcttccgcaaattccactatctgaccttcctgccctccacagatgatttctatgactgtgaggtggatcactggggtttggaggagcctctgcggaagcactgggagtttgaagagaaaaccctcctcccagaaactaaagagtgtgatgccacgttgaccgagaaaagctttgaaacagatatgaacctaaactttcaaaacctgtcagttatgggactccgaatcctcctgctgaaagtagcgggatttaacctgctcatgacgctgaggctgtggtccagttgaggatccgcta

The MCC:I-E^k^β.TCRβ chimeric CRM^pMHCII^^β^ subunit was subcloned into the “pP2-mEGFP” puromycin-resistance MSCV vector via 5’XhoI and 3’NotI. This resulted in the CRM^pMHCII^^β^ subunit being cloned in frame with a (GGGGS)x3 linker and mEGFP as previously reported for TCRβ. For the sequence below, the underlined nucleotides flank the MCC peptide-encoding sequence:

> aatctcgagcgccaccatggtgtggctccccagagttccctgtgtggcagctgtgatcctgttgctgacagtgctgagccctccagtggctttggtcagagactcc<u>ggatcc</u>gccaacgagagggccgacctgatcgcctacctgaagcaggccaccaag<u>gaattc</u>agatccggaggcggaggctccctggtgcctcggggctccggaggcggaggctccgtcgacagaccatggtttttggaatactgtaaatctgagtgtcatttctacaacgggacgcagcgcgtgcggcttctggtaagatacttctacaacctggaggagaacctgcgcttcgacagcgacgtgggcgagttccgcgcggtgaccgagctggggcggccagacgccgagaactggaacagccagccggagttcctggagcaaaagcgggccgaggtggacacggtgtgcagacacaactatgagatcttcgataacttccttgtgccgcggagagttgagcctacggtgactgtgtaccccacaaagacgcagcccctggaacaccacaacctcctggtctgctctgtgagtgacttctaccctggcaacattgaagtcagatggttccggaatggcaaggaggagaaaacaggaattgtgtccacgggcctggtccgaaatggagactggaccttccagacactggtgatgctggagacggttcctcagagtggagaggtttacacctgccaggtggagcatcccagcctgaccgaccctgtcacggtcgagtggaaagcacagtccacatctgcacagaacaagtgtggaatcactagtgcatcctatcatcagggggttctgtctgcaaccatcctctatgagatcctactggggaaggccaccctatatgctgtgctggtcagtggcctagtgctgatggccatggtcaagaaaaaaaattccgcggccgcatgatgagatctgagctccatagaggcg

The Hb:I-E^k^β.TCRβ chimeric CRM^pMHCII^^β^ subunit was generated identically to that described above for the MCC:I-E^k^β.TCRβ chimeric CRM^pMHCII^^β^ subunit. The nucleotides that follow encode the Hb peptide as well as the underlined nucleotides that flank the peptide-encoding sequence:

<u>ggatcc</u>ggcaagaaggtgatcaccgccttcaacgagggcctgaag<u>gaattc</u>

The following sequence for the I-A^d^α.TCRα chimeric CRM^pMHCII^*^α^* subunit was subcloned into the pMSCV-ires-CFP II (gift from Dario Vignali, Addgene plasmid # 52109) via 5’EcoRI and 3’XhoI:

> gaattccgccaccatgccgtgcagcagagctctgattctgggggtcctcgccctgaacaccatgctcagcctctgcggaggtgaagacgacattgaggccgaccacgtaggcttctatggtacaactgtttatcagtctcctggagacattggccagtacacacatgaatttgatggtgatgagttgttctatgtggacttggataagaagaaaactgtctggaggcttcctgagtttggccaattgatactctttgagccccaaggtggactgcaaaacatagctgcagaaaaacacaacttgggaatcttgactaagaggtcaaatttcaccccagctaccaatgaggctcctcaagcgactgtgttccccaagtcccctgtgctgctgggtcagcccaacacccttatctgctttgtggacaacatcttcccacctgtgatcaacatcacatggctcagaaatagcaagtcagtcacagacggcgtttatgagaccagcttcctcgtcaaccgtgaccattccttccacaagctgtcttatctcaccttcatcccttctgatgatgacatttatgactgcaaggtggagcactggggcctggaggagccggttctgaaacactgggaacctgagattccagcccccatgtcagagctgacagaaactgtgtgtgatgccacgttgaccgagaaaagctttgaaacagatatgaacctaaactttcaaaacctgtcagttatgggactccgaatcctcctgctgaaagtagcgggatttaacctgctcatgacgctgaggctgtggtccagttgactcgag

The RLGL-WE14:I-A^g7^β.TCRβ chimeric CRM^pMHCIIβ^ subunit was subcloned into the “pP2-mEGFP” puromycin-resistance vector via 5’XhoI and 3’NotI. This resulted in the CRM^pMHCIIβ^ subunit being cloned in frame with a (GGGGS)x3 linker and mEGFP as previously reported for TCRβ. For the sequence below, the underlined nucleotides flank the RLGL-WE14 peptide-encoding sequence:

> cacgcaagcttctcgagcgccaccatggctctgcagatccccagcctcctcctctcggctgctgtggtggtgctgatggtgctgagcagcccagggactgagggcggagactcc<u>gcggatccc</u>cgcttgggcttgtggagtaggatggaccaattagccaaggaattgactgcggag<u>gtcgac</u>ggaggtggcgggtcactagtgccccgaggaagtggaggtggagggtctccagggactgagggcggagactccgaaaggcatttcgtgcaccagttcaagggcgagtgctacttcaccaacgggacgcagcgcatacggctcgtgaccagatacatctacaaccgggaggagtacctgcgcttcgacagcgacgtgggcgagtaccgcgcggtgaccgagctggggcggcactcagccgagtactacaataagcagtacctggagcgaacgcgggccgagctggacacggcgtgcagacacaactacgaggagacggaggtccccacctccctgcggcggcttgaacagcccaatgtcgccatctccctgtccaggacagaggccctcaaccaccacaacactctggtctgttcggtgacagatttctacccagccaagatcaaagtgcgctggttcaggaatggccaggaggagacagtgggggtctcatccacacagcttattaggaatggggactggaccttccaggtcctggtcatgctggagatgacccctcatcagggagaggtctacacctgccatgtggagcatcccagcctgaagagccccatcactgtggagtggagggcacagtccgagtctgcccggagctgtggaatcactagtgcatcctatcatcagggggttctgtctgcaaccatcctctatgagatcctactggggaaggccaccctatatgctgtgctggtcagtggcctagtgctgatggccatggtcaagaaaaaaaattccgcggccgcatga

The genes encoding the GPI:I-A^g7^β.TCRβ, InsB9-23:I-A^g7^β.TCRβ, HIP2.5:I-A^g7^β.TCRβ, and HIP6.9:I-A^g7^β.TCRβ chimeric CRM^pMHCIIβ^ subunits were generated identically to that described above for the RLGL-WE14:I-A^g7^β.TCRβ chimeric CRM^pMHCIIβ^ subunit. The nucleotide sequences that follow encode the distinct peptides. The underlined nucleotides flank the peptide-encoding sequence:

GPI: <u>gcggatccc</u>ttatctattgcgcttcacgtgggcttcgatcactttgaa<u>gtcgac</u>
INSB 9-23: <u>gcggatccc</u>tcacatctagttgaagcgctatatctagtttgcggagaaagaggc<u>gtcgac</u>
HIP2.5: gcggatcccggcgacctgcagactctggccctgtggagcaggatggacgtcgac
HIP6.9: <u>gcggatccc</u>ggcgacctgcagactctggccctgaacgccgccagggac<u>gtcgac</u>

### Retroviral production

Retroviruses used for transduction of 58α^−^β^−^ cells, M12 cells, and B10.A-derived CTLs were produced as previously described^73^. For transduction of NOD-derived CTLs, stable Phoenix-eco cell lines were generated using amphotrophic retroviruses produced in 293T cells. 293T cells were transiently transfected with the retroviral pP2 MSCV-puro vector encoding the CRM^pMHCII^^β^ or ScoR constructs, or pMSCV-ires-CFP II (gift from Dario Vignali, Addgene plasmid # 52109) containing the CRM^pMHCII^*^α^* constructs, in addition to the packaging plasmid, pUMVC (gift from Bob Weinberg, Addgene plasmid # 8449)^74^, and the VSV-G envelope-expressing plasmid, pMD2.G (gift from Didier Trono, Addgene plasmid # 12259). Amphotrophic retroviral supernatant was collected 72 hr after transfection and used for the infection of Phoenix-eco cells (ATCC) by using Retronectin (Clontech). Transduced Phoenix-eco cells were sorted using either GFP, CFP, or CD80, as appropriate, and were used as stable producers of retroviruses for transduction of CTLs.

#### Cell lines and Mice

58α^−^β^−^ T cell hybridoma and M12 B cell lymphoma cell lines were cultured in RPMI 1640 (Gibco) supplemented with 5% FBS (Atlanta Biologicals), L-glutamine Pen-Strep solution (Hyclone), and 50mM 2-mercaptoethanol^26,27^. B10.A-H2 <A> H2-T18 <A>/SgSnJ (B10.A) mice were purchased from Jackson Laboratory and 5c.c7 TCR Tg × Rag2^−/−^ [B10.A-*Rag2^tm1Fwa^ H2-T18^a^* Tg (Tcra5CC7,Tcrb5CC7)lwep] mice were purchased from Taconic. They were maintained under specific pathogen-free conditions in the animal facility at the University of Arizona. NOD/ShiLtJ (NOD) mice, NOD.CB17-Prkdc ^scid^/J (NOD SCID) mice and NOD.Cg-Tg (TCRα^BDC2.5^,TCRβ^BDC2.5^)1Doi/DoiJ (NOD.BDC2.5) mice were purchased from Jackson Laboratory. NOD.mRaspberry (mRasp) transgenic mice were a gift from Dr. Jason Gaglia. NOD.BDC2.5.mRasp transgenic mice were generated by interbreeding of NOD.BDC2.5 TCR transgenic mice with NOD.mRasp transgenic mice. NOD.BDC2.5, NOD.mRasp and NOD.BDC2.5.mRasp transgenic mice were bred at the Joslin Diabetes Center Animal Facility (Boston, MA). Mice were 0-14 weeks-old at the initiation of experiments. All strains were maintained under specific pathogen-free conditions at the Joslin Diabetes Center Animal Facility. All experiments involving animals were conducted under guidelines and approval by the University of Arizona and Joslin Institutional Animal Care and Use Committees.

#### Antibodies and reagents for flow cytometry and cell sorting

The following antibodies were purchased from Biolegend: PE/Cy7 conjugated anti-mouse CD4 (GK1.5), CD8α (53-6.7), CD11b (M1/70), CD11b (N418), Gr-1 (56-8C5), B220 (RA3-6 B2), CD25 (PC.61) and Granzyme B (QA16A02) antibodies; APC/Cy7 conjugated anti-mouse CD3 (17-A2), CD8α (53-6.7) and IFN*γ* (XMG1.2) antibodies; APC conjugated anti-mouse CD8α (53-6.7), CD62L (MEL-14) and I-A^d^ (39-10-8) antibodies; PE conjugated anti-mouse CD4 (GK1.5), CD80 (16-10A1), Perforin (S16009B) and FasL (NOK-1) antibodies; PerCP/Cy5.5 conjugated anti-mouse CD4 (GK1.5) and Fas (SA367H8); Brilliant Violet (BV) 711 conjugated anti-mouse CD44 (IM7) antibody; BV 785 conjugated anti-mouse CD11b (M1/70); and, biotin conjugated anti-mouse CD3ε (145-2C11). The following antibodies and reagents were obtained from BD Biosciences: BV605 streptavidin; and, FITC, PE, and Biotin conjugated anti-Vβ4 T-cell receptor antibodies. The following antibodies were purchased from eBiosciences: PE conjugated anti-mouse MHCII (M5/114.15.2); PE-Cy7 conjugated anti-mouse CD3ε (145-2C11); and APC conjugated anti-mouse CD80 (16-10A1).

#### Generation of ^5M^CAR-CTL

Splenocytes from 5-10-week-old B10.A and NOD mice were stained with PE/Cy7 conjugated anti-mouse CD4 antibody. Negative selection was performed using anti-Cy7 microbeads (Miltenyi Biotech) according to the manufacturer’s protocol. Unbound cells were collected and cultured for 24 hours in the presence of 1 μg/ml anti-mouse CD3ε (145-2C11, BioLegend) and 0.5 μg/ml CD28 (37.51, eBioscience) in RPMI (Gibco) supplemented with 10% heat-inactivated fetal bovine serum (FBS; Gemini), 100 unit/ml penicillin and streptomycin (HyClone), 1 mM sodium pyruvate (Gibco) and 2 mM L-Glutamine (Gibco). ^5M^CAR retroviral supernatants: I-Ag^7^α-TCRα, peptide:I-A^g7^β-TCRβ and CD80-LCK, were mixed at a 1:1:1 ratio and bound to retronectin (Clontech)-coated plates according to the manufacturer’s protocol. The activated cells were added to the virus-bound plates after removing dead cells using Histopaque-1119 (Sigma-Aldrich), and centrifuged at 3200 × g for 90 min at 32°C. Murine IL-2 (50 unit/ml, Peprotech) was added immediately after transduction. Puromycin (2 μg/ml, Sigma-Aldrich) was added 36 hours after transduction for selection of ^5M^CAR expressing cells. 4-6 days after transduction, dead cells were removed using Histopaque-1119, and live cells were used for experiments. Typically, 85% of cells in the cultures expressed the transduced ^5M^CARs, as measured by flow cytometry.

#### Flow cytometry

Cells were stained with monoclonal antibodies, incubated on ice in staining buffer for 20 minutes, and washed. Some experiments utilized secondary staining with conjugated streptavidin. For flow cytometry of live cells, after the final wash, cells were resuspended in staining buffer with DAPI (1 μg/ml) or PI (100 ng/ml) to enable exclusion of dead cells. For intracellular IFN*γ*, Granzyme B or Perforin staining, 4 μM monensin (Sigma-Aldrich) was included in the final 3 hr of culture. Immediately following culture, cells were stained with Fixable Viability Dye eFluor 506 (eBioscience) to permanently stain dead cells. Surface molecules were stained, and then cells were fixed with 4% PFA at room temperature for 10 minutes and permeabilized with intracellular staining perm wash buffer (BioLegend), and stained with the appropriate intracellular antibodies. Analysis of stained cells was performed on an LSR Fortessa (BD Biosciences), and sorting was performed on a Cytomation MoFlo or a FACS Aria (BD Biosciences) at the Joslin Diabetes Center Flow Cytometry Core. Flow cytometry data were analyzed with FlowJo Version 10 software (Tree Star).

#### Flow-based fluorophore-linked immunosorbent assay (FFLISA)

6.0 µm streptavidin-coated polystyrene microspheres (Polysciences) were further coated with biotinylated anti-mouse CD3ε, washed, and incubated with 0.5 ml of DDM (1% *n*-dodecyl-b-D-maltoside) lysates from 5×10^6 5M^CAR 58α^−^β^−^ cells at 4°C for 1 hour. After washing, beads were probed with PE conjugated anti-mouse CD3*ζ* (clone G3, Santa Cruz Biotechnology) and analyzed by flow cytometry^14^.

#### ELISA

All reagents for detection of IL-2 were purchased from Biolegend. Supernatants were collected after 16 hours of coculture of ^5M^CAR 58α^−^β^−^ cells and M12 target cells. Anti-mouse IL-2 clone JES6-1A12 was used as a capture antibody and clone JES6-5H4 was used as a secondary antibody. Streptavidin-HRP and TMB substrate were used for detection.

#### Lymphocyte preparation

Lymphocytes were obtained by forcing spleens, pancreatic lymph nodes (pLNs) or non-draining LNs through 70 μm mesh. Red blood cells were lysed using ACK lysis buffer. Single cells from the pancreas were isolated by enzymic digestion with collagenase P. 1 mg/ml collagenase P (Roche) was directly injected into the pancreas. The pancreas was digested at 37°C for 8 minutes, blocked with PBS containing 10% FBS, and minced into small pieces on ice, dispersed by pipetting, filtered through a 70 μm mesh, wash with 10% FBS in PBS. Single mononuclear cells were purified by density gradient centrifugation using Histopaque 1.077 (Sigma-Aldrich).

#### *In vitro* killing, IFNγ production, and proliferation

For purification of CD4^+^ T cells, splenocytes from NOD.mRasp or NOD.BDC2.5.mRasp transgenic mice were stained with PE/Cy7 conjugated anti-mouse CD8α, CD11b, CD11c, B220 and Gr-1, and negatively selected using anti-Cy7 microbeads (Milttenyi Biotech) according to the manufactures’ protocol. The unbound CD4^+^ T cells were collected and used as target T cells. CD4^+^ T cell targets from 5c.c7 TCR Tg mice were not purified. Cells were co-cultured in 96-well U bottom plates. For killing assays, 1 × 10^5^ target T cells were cultured alone, or with ^5M^CAR-CTLs at different ratios for 16 hours. Target CD4 T^+^ cells were identified as Vβ3^+^CD4^+^GFP^−^CD8^−^ for 5c.c7 T cell targets and DAPI^−^ mRasp^+^CD4^+^GFP^−^CD8^−^ cells for BDC2.5 T cell targets. The numbers of target CD4^+^ T cells that remained after 16hrs culture were then enumerated by flow cytometry. The number of target cells that remained in the absence of ^5M^CAR-CTLs was used to establish a no-killing baseline that was then used to calculate the percent killing in the ^5M^CAR-CTL samples. For the analysis of IFNγ production, 1 × 10^4 5M^CAR-CTLs were cultured with 1 × 10^5^ target T cells for 6 hours. For the analysis of proliferating ^5M^CAR-CTL, 1 × 10^4 5M^CAR-CTLs labeled with 2 μM CellTrace Violet (Invitrogen) were cultured with 1 × 10^5^ NOD.BDC2.5.mRasp target CD4^+^ T cells for 3-4 days. ^5M^CAR-CTLs were identified as GFP^+^CD8^+^mRasp^−^CD4^−^ cells. Absolute cell numbers were calculated using CountBright Absolute Counting Beads (Molecular Probes) on flow cytometry, or were counted directly by running the entire sample on the flow cytometer.

#### *In vivo* killing assay

CD4^+^ T cells were purified from the spleens of NOD or NOD.BDC2.5.mRasp transgenic mice by negative selection as described above and were used as target T cells. NOD CD4^+^ T cells were labeled with 10 μM CellTrace Violet. 5-week-old female NOD recipient mice were retro-orbitally transplanted with GPI:I-A^g7^ or RLGL-WE14:I-A^g7 5M^CAR-CTLs (7 × 10^6^ cells per mouse), and 12 hours later were transplanted with a 1:1 mix of CellTrace Violet labeled NOD CD4^+^ T cells and NOD.BDC2.5.mRasp CD4^+^ T cells (each 3.5 × 10^6^ target cells per mouse). 5.5 hours later the ratio of mRasp^+^ BDC2.5 T cells to CellTrace Violet-labeled NOD T cells in the spleen of each recipient was determined by flow cytometry.

#### Adoptive transfer

For the experiments using the NOD-SCID T1D model, mRasp^+^CD4^+^CD8^−^CD25^−^CD62L^+^Vβ4^+^ cells were obtained from the spleens of NOD.BDC2.5.mRasp transgenic mice and intravenously transplanted into NOD.SCID recipients (3 × 10^5^ cells per recipient). 1 or 7 days later mice were received GPI:I-A^g7^ or RLGL-WE14:I-A^g7 5M^CAR-CTLs (6 × 10^6^ cells per mouse). The mice were screened daily for body weight and glycosuria and/or blood glucose level. Mice were euthanized one day after diabetes onset, defined as two consecutive positive readings. Diabetes-free recipients were euthanized at the end point of experiment (day 21 or day 36). Target T cells were identified as DAPI^−^mRasp^+^CD3^+^GFP^−^ or DAPI^−^mRasp^+^CD3^+^CD11b^−^ GFP^−^ cells, and ^5M^CAR-CTLs were identified as DAPI^−^GFP^+^CD3^+^mRasp^−^ or DAPI^−^ GFP^+^CD3^+^CD11b^−^mRasp^−^ cells. For the experiments using NOD mice as recipients, 3-5 day old NOD mice were transplanted by facial vein injection with GPI:I-A^g7 5M^CAR-CTLs (1 × 10^6^ cells per mouse) or a mixture of ^5M^CAR-CTLs (INSB:I-A^g7^, HIP2.5:I-A^g7^, HIP6.9:I-A^g7^ and RLGL-WE14:I-A^g7^, each 2.5 × 10^5^ cells, totaling 1 × 10^6^ cells per mouse) and were screened weekly for glycosuria staring at 8 weeks of age. Diabetes was defined as two consecutive weekly readings positive for glycosuria.

#### Histology

Pancreases were harvested, embedded in OCT and frozen in isopentane. Pancreas sections were cut and stained with hematoxylin-eosin. Photos were taken by an Olympus VS120 slide scanner (Olympus) with x20 or x40 objective at Harvard Medical School Neurobiology Imaging Facility. Images were acquired on an Olympus Olyvia at 2X, 4X or 20X magnification. Data was analyzed on GIMP software (The GNU image manipulation program, ver.2.8).

#### Statistical analysis

Statistical analysis was performed with GraphPad Prism software (La Jolla, CA) and with SAS software (v.9.4, Carry, NC). Results are expressed as mean ± SD. Statistical significance of differences between the groups was determined as indicated in the figure legends for each experiment. Cox proportional hazards models were performed with ties in the failure time handled with exact conditional probabilities. P< 0.05 was considered statistically significant.

## Acknowledgements

This work was supported by The University of Arizona College of Medicine (MSK), the BIO5 Institute (MSK), National Institutes of Health/National Institute of Allergy and Infections Diseases Grant R01AI101053 (MSK), the Pew Scholars Program in the Biomedical Sciences (MSK), charitable donations to the UA Foundation (MSK), the Cancer Center Support Grant CCSG-CA 023074 for flow cytometry (MSK), the Fleisher Family Foundation (TS), the Alexander and Margaret Stewart Trust (TS), the Iacocca Family Foundation (SK, MAT), the Swedish Society of Medicine and the Swedish Society of Medical Research (MAT). Statistical analysis (MAN) was conducted with support from Harvard Catalyst | The Harvard Clinical and Translational Science Center (National Center for Advancing Translational Sciences, National Institutes of Health Award UL 1TR002541). The JDC Flow Cytometry Core is supported by a National Institute of Health grants (P30-DK-036836 and S10OD021740). We thank Amy Wagers for additional support and critical feedback. We thank David Duron, Karen Hernandez, and Lacey Orsini for technical assistance with generation of constructs. The authors also thank Deepta Bhattacharya, Alfred Bothwell, Michael Worobey, and Joonsoo Kang for critically reading the manuscript.

## Author’s Contributions

M.S.K. and T.S. conceived of the project and directed the research. M.S.K. conceived of and designed the ^5M^CARs. The manuscript was written by S.K., T.S., and M.S.K. All authors contributed to data analysis, discussions, editing, reading and approval of the manuscript. H.L.P., N.R.D., and M.S.L. generated and performed *in vitro* experiments with the ^5M^CAR-58α^−^β^−^ cells and B10.A-derived ^5M^CAR-CTLs, while S.K. and M.A.T. generated and performed *in vitro* experiments with NOD-derived ^5M^CAR-CTLs. S.K. and M.A.T. performed experiments in NOD-SCID mice while S.K. and A.K. performed long-term experiments in NOD mice. M.A.N. performed statistical analysis.

## Supplementary Figure Legends

**Supplementary Figure 1 (Corresponding to Fig 1):**
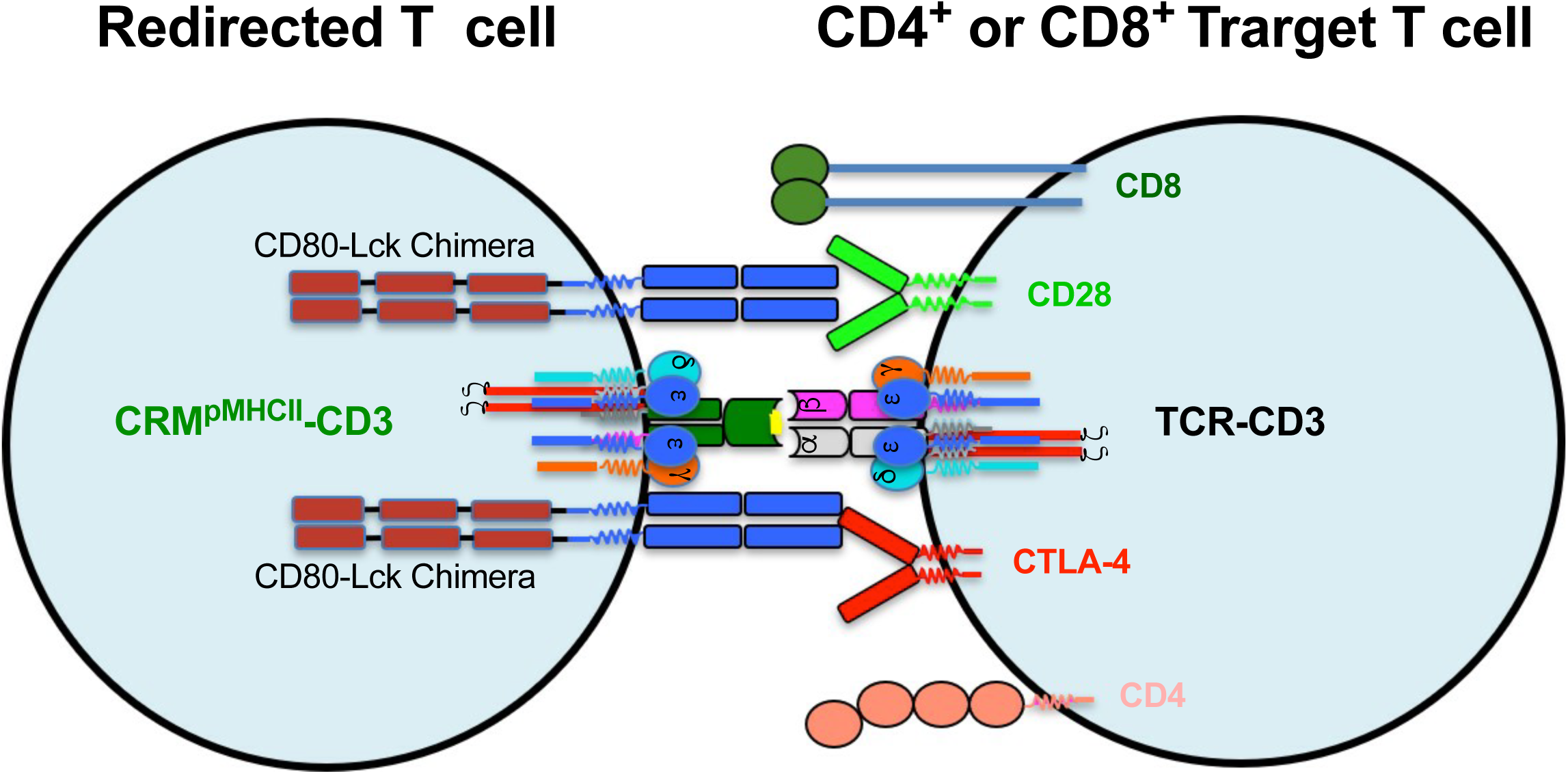
Design of ^5M^CAR system. **a**, Drawing of proposed interaction between ^5M^CAR-CTLs and pathogenic CD4^+^ T cells.

**Supplementary Figure 2 (Corresponding to Fig 2):**
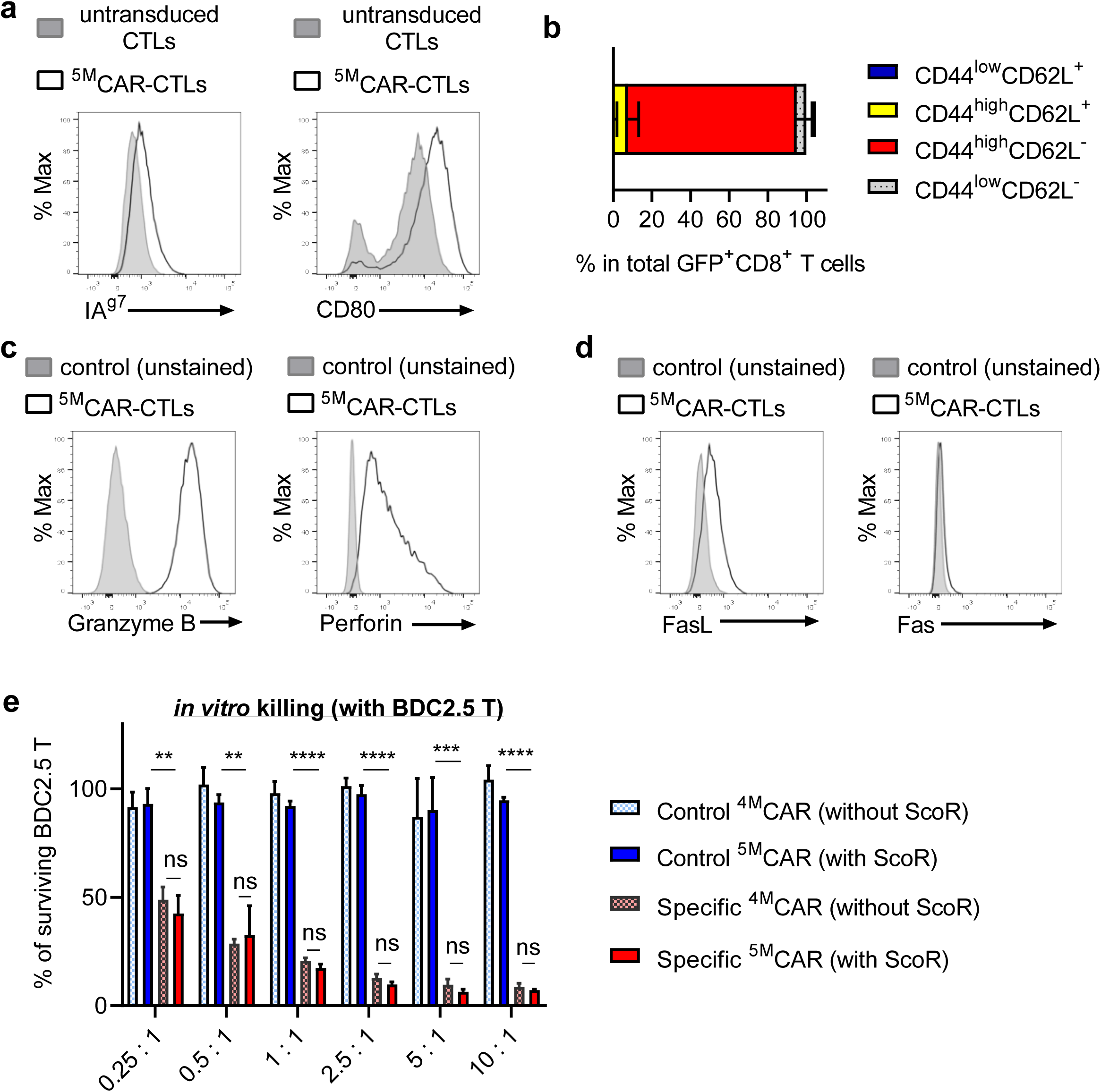
^5M^CAR-CTL phenotype and killing. **a**, Representative histograms showing I-A^g7^ (left) and CD80 (right) expression on ^5M^CAR-CTLs (solid line) compared with untransduced CD8^+^ CTLs (shaded histogram). Data represent at least three independent experiments. **b**, CD44 and CD62L expression of ^5M^CAR-CTLs were measured. Data shows the proportion of each ^5M^CAR-CTL subset as mean ± SD of triplicates from three independent experiments. **c**, Representative histograms of Granzyme B (left) and Perforin (right) expression by ^5M^CAR-CTLs. **d**, Representative histograms of FasL (left) and Fas (right) expression by ^5M^CAR-CTLs. **a-d**, RLGL-WE14:I-A^g7 5M^CAR-CTLs 4 days post-transduction were stained. Histograms were gated on GFP^+^CD8^+^ cells. **e**, Normalized frequency of surviving target BDC2.5 T cells after a 16 hr coculture with either ^4M^CAR-CTLs (without ScoR) or ^5M^CAR-CTLs (with ScoR) at several E:T ratios. Control CARs were GPI:IA^g7^ and Specific CARs were (RLGL-WE14:I-A^g7^). Surviving target cells were defined as DAPI^−^ mRasp^+^CD4^+^GFP^−^CD8^−^. **p < 0.01, ***p < 0.001, ****p < 0.0001 by unpaired, two-tailed *t*-test. Columns are shown as mean ± SD of triplicates. The graph is representative of 2 independent experiments.

**Supplementary Figure 3 (Corresponding to Fig 3):**
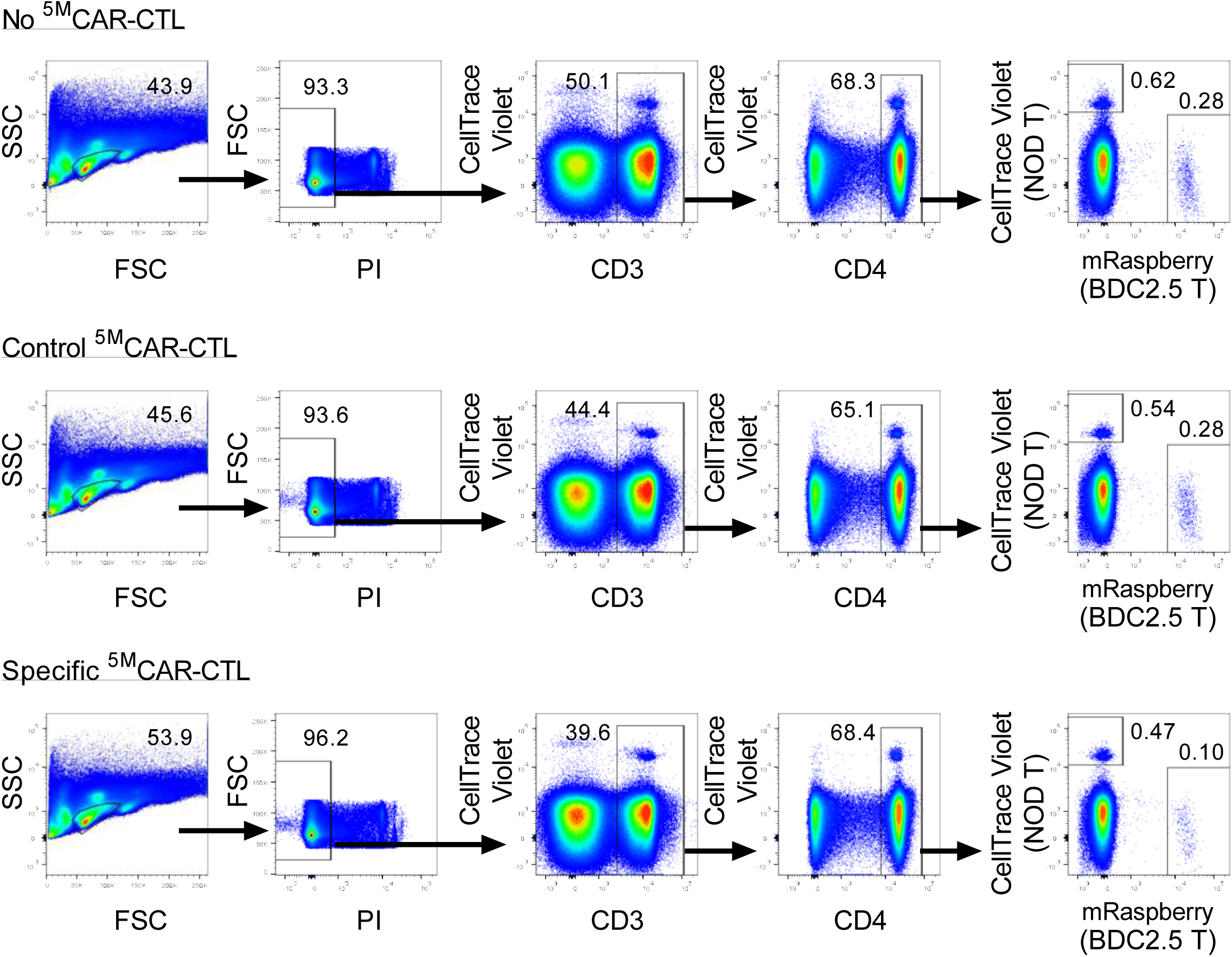
Gating scheme for *in vivo* killing assay. Flow cytometry of target T cells and ^5M^CAR-CTLs in NOD mice. Representative FACS plots show the gating strategy to identify mRasp^+^ CD4^+^ BDC2.5 target T cells and CellTrace Violet^+^ NOD control T cells for the data shown in Fig. 3.

**Supplementary Figure 4 (Corresponding to Fig 4):**
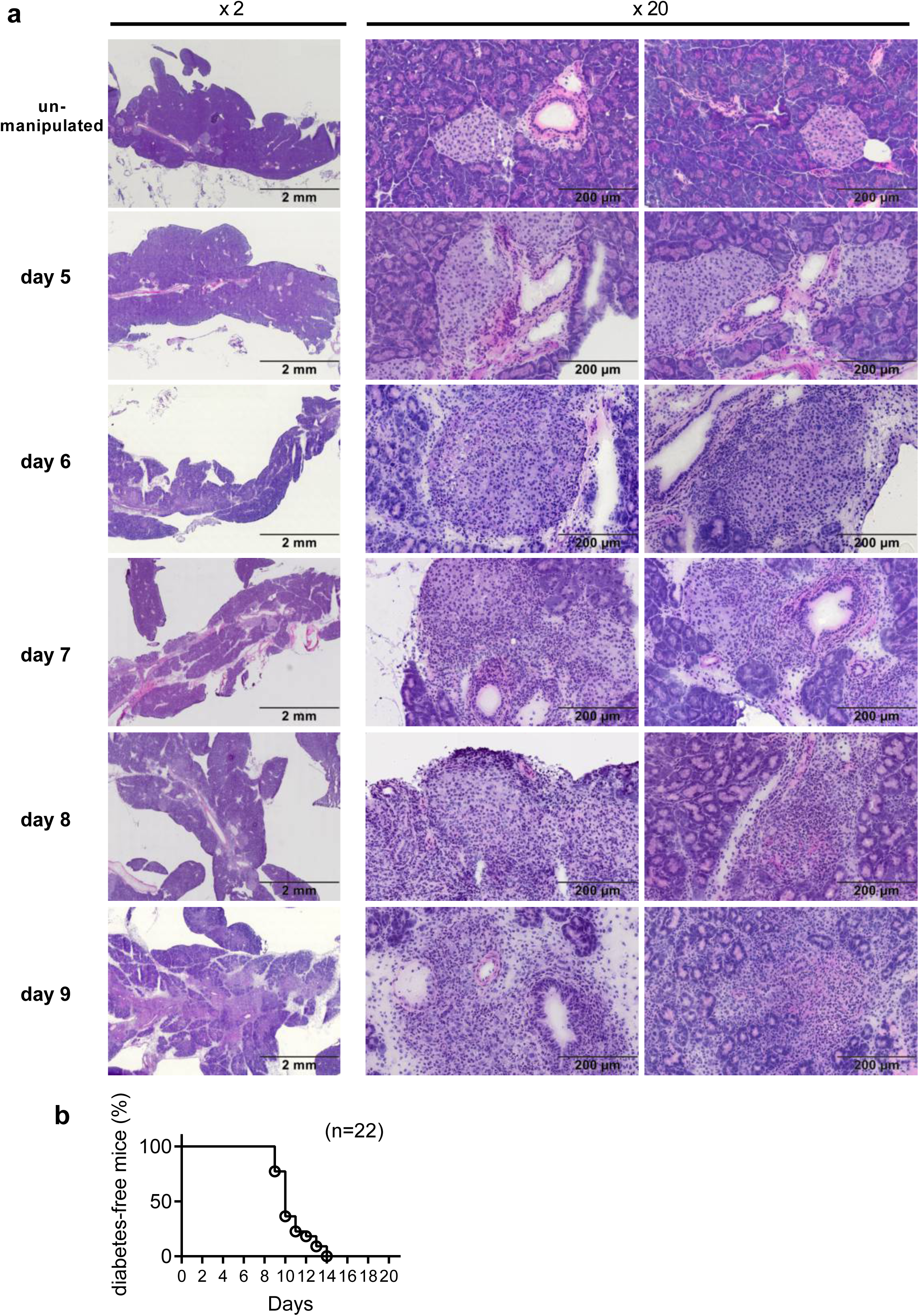
Time course of insulitis and diabetes development in NOD-SCID mice after BDC2.5 CD4^+^ T cell transfer. BDC2.5 CD4^+^ T cells were adoptively transferred into NOD-SCID mice on day 0 and euthanized on day 5, 6, 7, 8 or 9. **a**, Pancreases of representative mice from each group are shown stained with hematoxylin-eosin (Magnification: Left column 2X, Right columns 20X). Islets from an unmanipulated mouse show no cell infiltration and are distinctly separated from adjacent acinar tissue (top row). Immune cell infiltration of islets was observed at day 5. Infiltrating cells localized to the ductal pole of the islet and connective tissue. At day 6 and 7, infiltrating cells have surrounded and invaded into islets, which show compromised borders. By day 8 and 9, the architecture of islets has been destroyed by massive cell infiltration. All mice were diabetes-free at each time point. **b**, Kaplan-Meier curve shows the time course of diabetes development in mice transplanted with 300,000 BDC2.5 T cells on day 0. Data are combined from four independent experiments.

**Supplementary Figure 5 (Corresponding to Fig 5):**
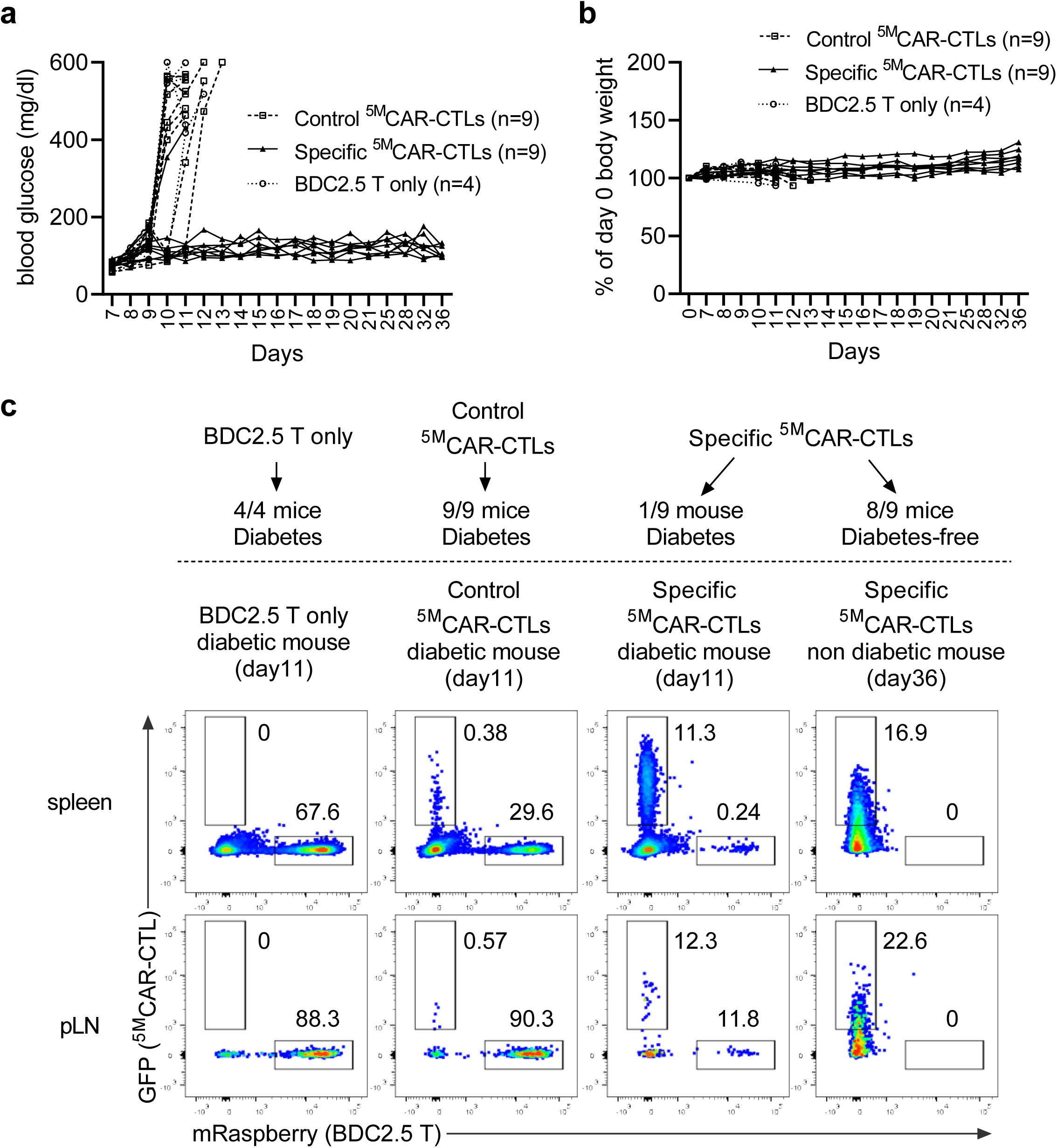
Mice infused with specific ^5M^CAR-CTLs remain normoglycemic, maintain body weight, and eliminate target T cells. BDC2.5 CD4^+^ T cells were adoptively transferred into NOD SCID mice on day 0. On day 7 the mice were treated with either control (GPI:I-A^g7^) ^5M^CAR-CTLs, specific (RLGL-WE14:I-A^g7^) ^5M^CAR-CTLs, or left untreated (BDC2.5 only). **a**, Blood glucose levels and **b**, weight changes of recipients are shown. Weight change was calculated on the basis of the weight at day 0. **a**,**b**, Data shows the combined results from two independent experiments. **c**, Representative dot plots show frequencies of targets and ^5M^CAR-CTLs in spleens or pLNs of treated and untreated mice. Diabetic mice were euthanized and analyzed one day after they developed diabetes (day 11 for mice represented on plots). Diabetes-free mice were analyzed at the experimental endpoint, day 36. Data represent two independent experiments. Numbers of mice/group is indicated in the figure.

**Supplementary Figure 6 (Corresponding to Fig 6):**
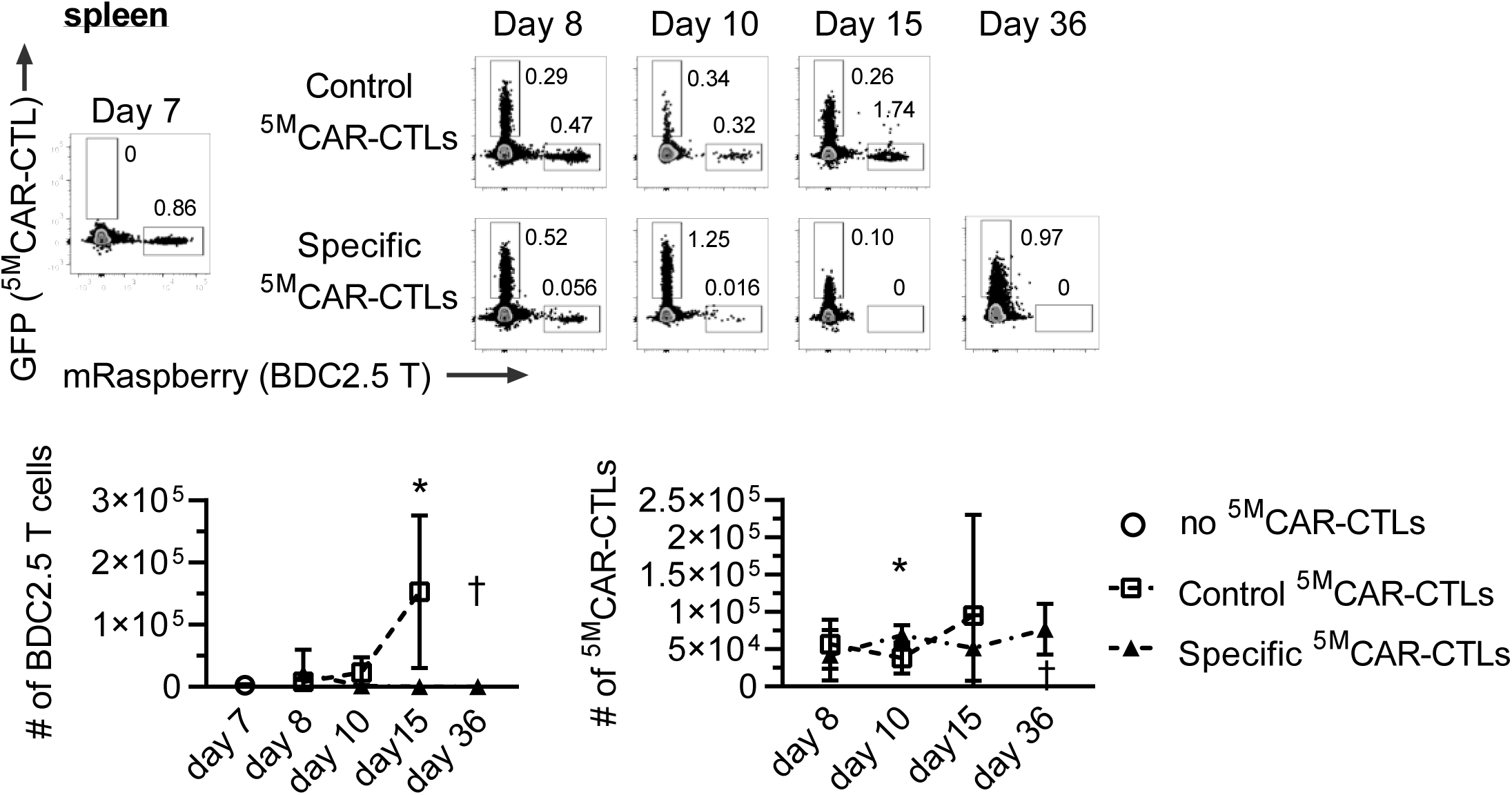
Enumeration of ^5M^CAR-CTLs and BDC2.5 CD4^+^ T cells in the spleen. NOD-SCID mice received BDC2.5 CD4^+^ T cells on day 0 and were either treated on day 7 with control (GPI:I-A^g7^) ^5M^CAR-CTLs, or specific (RLGL-WE14:I-A^g7^) ^5M^CAR-CTLs, or were euthanized pre-treatment. Treated groups were euthanized on day 8, 10, 15 or 36. Flow cytometry was used to determine the frequency and number of mRasp^+^ CD4^+^ BDC2.5 T cell target and GFP^+ 5M^CAR-CTLs in the spleens. Representative dot plots show frequencies of BDC2.5 CD4^+^ T cells and ^5M^CAR-CTLs (top panels, pre-gated on live, CD11b^−^ cells). Graphs show the number of BDC2.5 CD4^+^ T cells (left) or change in the number of ^5M^CAR-CTLs (right). Data show results combined from two independent experiments as mean ± SD. 3-6 mice were analyzed for each group and time point (*p < 0.05 by unpaired, two-tailed *t*-test between the control and specific ^5M^CAR-CTLs at each time point). † means no data.

**Supplementary Figure 7 (Corresponding to Fig 7):**
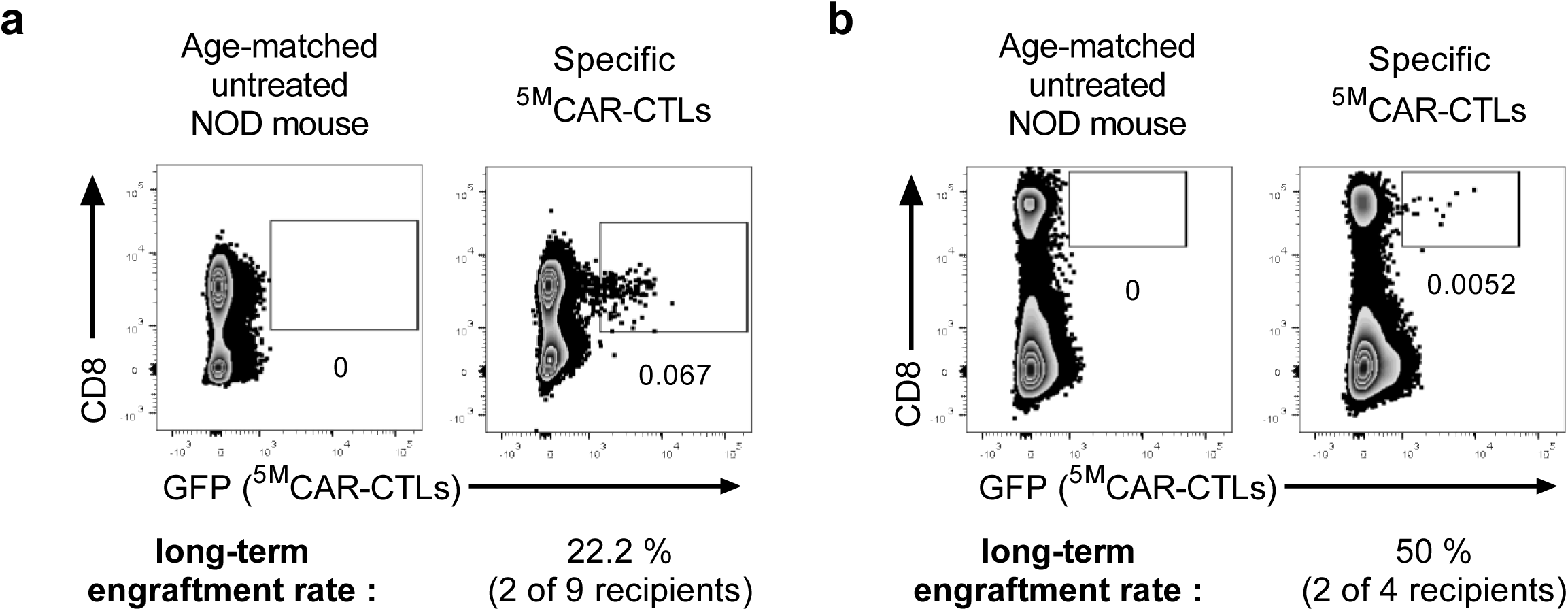
Long-term persistence of ^5M^CAR-CTLs in diabetes-free NOD mice. Newborn NOD mice were treated with control (GPI:I-A^g7^) ^5M^CAR-CTLs, a mixture of ^5M^CAR-CTLs (INSB:I-A^g7^, HIP2.5:I-A^g7^, HIP6.9:I-A^g7^ and RLGL-WE14:I-A^g7^), or left untreated and were monitored for glycosuria. Diabetes-free male and female mice were euthanized and lymphocytes were analyzed by flow cytometry at the conclusion of the experiment shown in Fig 7 (>315 days). **a,** Representative dot plots show the frequencies of long-term engrafted GFP^+^CD8^+ 5M^CAR-CTLs within the spleens plus the pLNs. **b**, The spleens and pLNs of boosted female recipients were harvested 2 days after BDC2.5 transfer and analyzed as a mixed sample. An age-matched unmanipulated female NOD mouse was used as a control. The proportion of mice with long-term engrafted ^5M^CAR-CTLs in each subgroup of diabetes-free recipients is shown.

